# Zinc deficiency induces spatially distinct responses in roots and impacts ZIP12-dependent zinc homeostasis in Arabidopsis

**DOI:** 10.1101/2025.06.26.661794

**Authors:** Noémie Thiébaut, Daniel Pergament Persson, Manon C. M. Sarthou, Pauline Stévenne, Bernard Bosman, Monique Carnol, Steven Fanara, Nathalie Verbruggen, Marc Hanikenne

**Author notes:** Corresponding author : Marc Hanikenne, InBioS – PhytoSystems, University of Liège, Quartier de la Vallée, 1, Chemin de la Vallée, 4 - Bât B22, B4000 Liège, Belgium.

## Abstract

How zinc (Zn) deficiency shapes root development remains unclear, with conflicting reports on its effect on primary root growth in *Arabidopsis thaliana* (Arabidopsis). The impact of Zn shortage on the root apical meristem (RAM) in particular has not been systematically explored. Using an integrative approach combining cell biology, transcriptomics, and ionomics, we dissected how Zn deficiency alters root zonation and function. We showed that Zn deficiency triggers a striking reorganization of the root tip (RT): the RAM size is reduced, yet meristematic activity and local Zn levels are preserved. This is accompanied by promoted cell elongation and differentiation. Transcriptome profiling revealed a distinct Zn deficiency response in the RAM-enriched RT compared to mature root tissues, with *ZIP12* emerging as the most strongly induced gene in the RT. Functional analysis of *zip12* mutants uncovered major defects in root growth, RAM structure, expression of Zn-responsive genes, and metal partitioning. Our work unveiled a new layer of root developmental plasticity under Zn deficiency and identified *ZIP12* as a central player in maintaining Zn homeostasis and root meristem function in *Arabidopsis*. These findings provide a framework to better understand how plants adapt root growth to fluctuating micronutrient availability.

## Introduction

Low zinc (Zn) availability in soil is one of the many challenges faced by plants in large natural and agricultural areas worldwide. The resulting Zn deficiency has severe consequences for human health, particularly in countries with a predominantly plant-based diet (Wessells & Brown, 2012; Assunção, 2022). Zn deficiency reduces plant growth as well as crop production and nutritional quality (Alloway, 2008; Thiébaut & Hanikenne, 2022; Lilay *et al*., 2024), and severely impacts the capacity of plants to respond to other stresses such as drought (Cakmak & Kutman, 2018; Noulas *et al*., 2018; Hassan *et al*., 2020), pathogen (Cakmak *et al*., 2000; Martos *et al*., 2016; Cabot *et al*., 2019) high light (Cakmak *et al*., 2000; Lilay *et al*., 2024), or exposure to toxic ions (Cakmak *et al*., 2000). Consequently, Zn deficiency exacerbates other global agricultural challenges. Indeed, it has pleiotropic effects on plants, ranging from imbalance in hormonal regulation, redox status or ion homeostasis, to reduced photosynthetic efficiency, increased light sensitivity and stunted growth (see Lilay *et al*., (2024) for a recent review).

Among dicots, the response to Zn deficiency is best described in the model plant *Arabidopsis thaliana* (Arabidopsis) (Lilay *et al*., 2019; Amini *et al*., 2022; Stanton *et al*., 2022; Assunção *et al*., 2022; Thiébaut & Hanikenne, 2022). Zn sensing and regulation are mediated by the bZIP19 and bZIP23 (basic leucine zipper) transcription factors, which trigger a massive induction of genes encoding zinc-regulated/iron-regulated-transporter-like proteins (ZIP) and nicotianamine synthases (NAS) under Zn deficiency (Assunção *et al*., 2010; Lilay *et al*., 2019, 2021; Assunção, 2022). Several ZIP transporters were shown to contribute to enhanced Zn uptake by roots, Zn radial transport in roots, and Zn root-to-shoot translocation (Talke *et al*., 2006; Lin *et al*., 2009; Ochoa Tufiño *et al*., 2025). Additionally, nicotianamine (NA) contributes to Zn root-to-shoot translocation by improving radial Zn transport in roots and Zn movement in the xylem (Clemens, 2019). Apart from Zn sensing in roots by bZIP19 and bZIP23, the Zn deficiency response is also regulated by yet-to-be identified shoot signalling (Sinclair *et al*., 2018). Moreover, Zn deficiency affects the homeostasis of other nutrients. For instance, the Zn deficiency response may lead to an increase of iron (Fe) concentration in plant tissues, caused by (i) the broad metal specificity of transporters, such as ZIPs, and (ii) the high affinity of ligands such as NA for several transition metals (Cakmak, 2000; Milner *et al*., 2013; Clemens, 2019; Thiébaut & Hanikenne, 2022).

Research and breeding efforts have mostly addressed Zn uptake and homeostasis mechanisms as a main focus to tackle Zn deficiency-related challenges (Garcia-Oliveira *et al*., 2018; Thiébaut & Hanikenne, 2022; Assunção, 2022; Robe & Barberon, 2023; Huizinga *et al*., 2025). The broader impact of Zn deficiency on other important plant traits has, however, been largely neglected, although they may be beneficial for coping with additional stresses in the context of global change (Lynch *et al*., 2021; Lynch, 2022; Lilay *et al*., 2024). Hence, reduced root growth is a hallmark phenotypic consequence of stress in plants. For instance, modifications in root system architecture have been proposed as promising breeding traits for adaptation to drier climate (Varshney *et al*., 2021; Lynch *et al*., 2021; Lynch, 2022). Similarly, identifying the determinants of root growth upon Zn deficiency could uncover mechanisms that could mitigate the effects of low Zn availability in soils.

Maintenance of root growth takes place at the root apex and involves regulation of both quiescence and meristematic activity in the root apical meristem (RAM). This includes mitosis, as well as the onset of endocycling (*i.e.* nuclear genome replication in the absence of mitosis), which is concomitant to the entry of the cells in the elongation and differentiation stages of root development (Bhosale *et al*., 2018). Zinc is fundamental to plant growth, as it is required as cofactor for ∼10% of the plant proteome (Ireland & Martin, 2019; Clemens, 2022), including for the structure and activity of several major transcription factor families (Ciftci-Yilmaz & Mittler, 2008; Phukan *et al*., 2016; Fedotova *et al*., 2017; Khoso *et al*., 2022) and other proteins of the core cell cycle machinery (Kipreos & Pagano, 2000; Bochman & Schwacha, 2009; Chanfreau, 2013; Clemens, 2022).

Whereas the impact of excess Zn (and other metals) on the cell cycle in the RAM and more widely on root growth has been described in details (Eekhout *et al*., 2017; van Dijk *et al*., 2022; Richtmann *et al*., 2025; Thiébaut *et al*., 2025), only a limited number of studies has examined which phases of the cell cycle are affected by Zn deficiency in plants and algae (Falchuk *et al*., 1975; Francis *et al*., 1995; Chesters & Petrie, 1999). Using synchronized BY-2 tobacco cells and the unicellular algae *Euglena gracilis,* these studies pointed antagonistically to either the S, G2-M, or G1 phases as limiting steps under Zn deficiency, while refuting Zn importance in other phases (Falchuk *et al*., 1975; Francis *et al*., 1995; Chesters & Petrie, 1999). However, recent and comprehensive studies in plants are lacking.

Here, we investigated the impact of Zn deficiency on root morphology and growth of the root apex in Arabidopsis. We showed that Zn deficiency reduced primary root growth and altered the size of the RAM, with a decrease in the number of dividing cells, accompanied by reduced cell elongation and differentiation. The deficiency did not, however, alter the cell cycle progression. Using complementary Zn element bioimaging and transcriptomics approaches, we identified distinct and different Zn deficiency responses in the RAM compared to differentiated roots, with notably a greater ability of the RAM to maintain cellular Zn levels. We further identified *ZIP12* as the most up-regulated ZIP transporter gene in the root tip (RT) upon Zn deficiency. Characterization of the growth and ionomic phenotypes of z*ip12* mutant plants revealed its key role in metal homeostasis upon severe Zn deficiency in Arabidopsis.

## Material and methods

Extended method descriptions are provided in the **Methods S1** text, as supporting information.

### Plant material, culture media and culture conditions

*Arabidopsis thaliana* (Col-0) was used in all experiments. The *ZIP12* (*AT5G62160*) mutants [*zip12-1* (SALK_137184), *zip12-2* (SALK_118705)], as well as the pCYCB1;2:CYCB1;2:GUS (N799897) and pCYCA3;1:CYCA3;1:GUS (N799893) lines were obtained from NASC (Nottingham Arabidopsis Stock Centre, UK) (**Fig. S1**, **Table S1**). The following lines were also used: pWOX5:GUS (Sarkar *et al*., 2007), pZIP4:GUS [BG0011, (Lin *et al*., 2016)], and *atm-1* (Garcia *et al*., 2003).

After surface sterilisation (Lindsey *et al*., 2017), seeds were sown directly on sterile petri plates with 50 mL of EDTA-washed agar Hoagland medium, which contained ZnSO_4_ in various concentrations (0 µM, 1 µM, 2 µM or 5 µM), as indicated. Growth conditions were as described in Thiébaut *et al*. (2025). Unless otherwise stated, all experiments were conducted in three independent replicates. The replication level of each experiment is detailed in figure legends.

### Root growth and RAM phenotyping

Primary root length of two-week-old plantlets was measured on images of scanned plates. Confocal microscopy images of propidium iodide-stained roots were used to measure the length of meristematic (Perilli & Sabatini, 2010) and elongation zones, as well as the size and number of RAM cortex cells. FIJI (https://imagej.net/Fiji) was used for all measurements.

### GUS staining and MUG assay

GUS staining and quantification of pZIP4:GUS, pWOX5:GUS, pCYCA3;1:CYCA3;1:GUS and pCYCB1;2:CYCB1;2:GUS were performed as previously described (Charlier *et al*., 2015; Thiébaut *et al*., 2025). The stained plantlets were then imaged with a Nikon SMZ1500 stereomicroscope equipped with a Nikon Digital Sight DS-5M camera (Nikon, Japan). Cell cycle synchronization was achieved using hydroxyurea, as described (Cools *et al*., 2010).

### RNA preparation

For RNA-Seq analysis, each root tip (RT) sample (∼2 mm length) was collected from the primary roots of ∼650 plantlets (Richtmann *et al*., 2025; Thiébaut *et al*., 2025), the remaining roots (RR) were pooled in a second associated sample. Total RNAs were extracted with the Maxwell® RSC Plant RNA Kit (Promega, USA) using a Maxwell® RSC robot (Promega, USA) and quality was controlled on a Bioanalyzer using a QC RNA kit (Agilent Technologies, USA) prior to RNA-Seq library preparation.

For Reverse Transcription-quantitative Polymerase Chain Reaction (RT-qPCR), samples were collected in biological triplicates. Each sample, ∼160 plantlets, was ground using a MM200 mixer mill (Retsch, Germany). Total RNA was extracted using the RNAeasy Microkit (Qiagen, Germany).

### RT-qPCR

cDNAs were prepared from 300 ng total RNA using Oligo(dT) with the RevertAid H Minus First Strand cDNA Synthesis Kit (Thermofischer, USA). RT-qPCR was conducted in technical triplicates for each 15x-diluted cDNA sample/primer combination with the Takyon Low Rox SYBR MasterMix dTTP Blue kit (Eurogentec, Belgium) on a QuantStudio5 thermocycler (Applied Biosystems, Thermofischer, USA) with the primers listed in **Table S1**. Primer pair efficiency was calculated using the LinReg PCR software (Ramakers *et al*., 2003) and expression was normalized with the expression of three housekeeping genes [*At5g60390-EF1alpha*, *At4g05320-UBQ10*, and *At1g58050,* (Nouet *et al*., 2015)] and relative to the control (2 µM Zn) using the qBase+ software (v.3.4.1, Biogazelle, Belgium).

### RNA-Seq, library preparation, sequencing, data analysis and representation

Library preparation (from 500 ng total RNA), quality control, quantification and RNA-Sequencing performed on a NovaSeq6000 were conducted at the GIGA Genomics platform of ULiège (Flowcell S4 V1.5, with 2x paired-end reads of 150 nucleotides). Data quality assessment and trimming, read mapping, counting and differential expression analysis were performed as described (Scheepers *et al*., 2020; Thiébaut *et al*., 2025).

A manually assembled list of cell cycle marker genes used for analysis is presented in **Table S2**. GO enrichment analysis for biological processes was conducted using the Panther Database (https://pantherdb.org).

Identification of the transcription factors was performed by comparing DEG lists to the *A. thaliana* Transcription Factor DataBase (AtTFDB) from the Arabidopsis Gene Regulatory Information Server (AGRIS, https://agris-knowledgebase.org/AtTFDB/).

### Mineral analysis

Shoots and roots of 160-180 two-week-old plantlets per sample were separated, then desorbed and washed (Scheepers *et al*., 2020; Thiébaut *et al*., 2025). Dried root and shoot samples were weighted, digested in HNO_3_ using a high-temperature digestion block, and analysed by ICP-OES (Inductively Coupled Plasma Optical Emission Spectroscopy, 5100 ICP-OES, Agilent Technologies, USA) (Nouet *et al*., 2015).

### Zinc imaging by Laser Ablation ICP-MS

Roots of two-week-old plantlets were harvested and two samples from the two regions of interest [*i.e.* the RAM (at ∼200 µm distance from the tip of the columella) and differentiated roots (at a distance from the root apex corresponding to 50-60 % of the total root length)] were prepared by encapsulation in melted paraffin and then frozen in a block of freezing media, as described (Thiébaut *et al*., 2025). Fourteen-µm thick sections were prepared (Persson *et al*., 2016), freeze-dried, photographed and analysed using a LA-ICP-MS system (Laser Ablation Inductively Coupled Plasma Mass Spectrometry).

## Results

### Zn deficiency impacts root growth

Compared to control growth conditions (1 µM Zn), both shoot and root growth was inhibited in Arabidopsis plantlets that were germinated and grown for 2 weeks on Zn-deprived solid agar medium (*i.e.* without any added Zn, 0 µM Zn) [**Fig. 1a, c-d,** (Thiébaut *et al*., 2025)]. The inhibition of primary root growth was accompanied by a reduction in size of the RAM, measured both in µm and in number of cortical cells present in the division zone (**Fig. 1b, e-f**). Moreover, the cortex cells in the division zone (delimited by the red arrows in **Fig. 1b**) were longer in Zn deficiency-exposed RAM (**Fig. 1b, g**).

**Figure 1.**
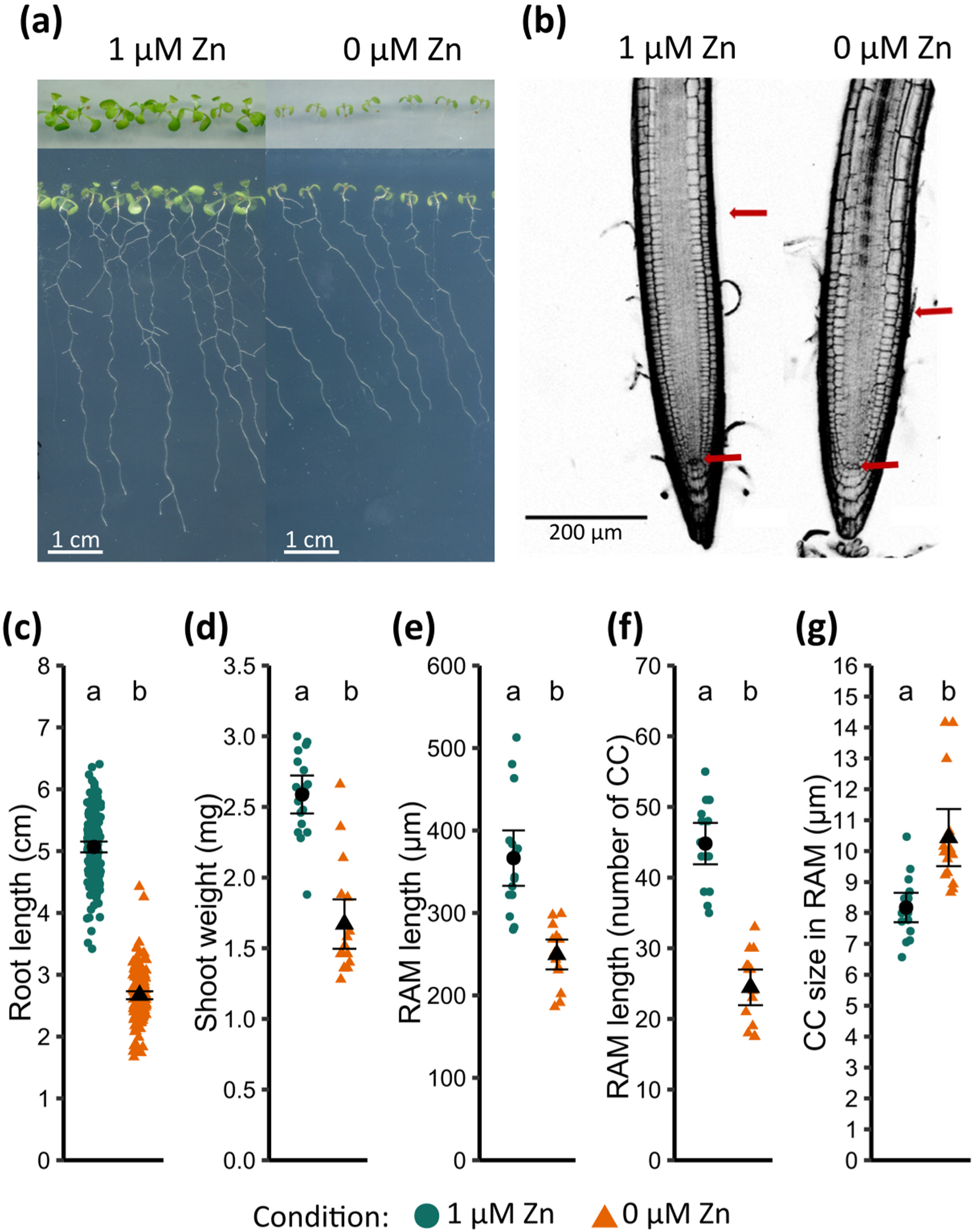
Primary root growth inhibition and impact on the root apical meristem upon Zn deficiency in Arabidopsis. Plantlets were grown for two weeks on EDTA-washed agar plates in control (1 µM Zn) or Zn deficiency (0 µM Zn) conditions. **a.** Representative pictures of the shoots (top) and roots (bottom). **b.** Representative images of root apical meristem (RAM). RAM size was measured between the two red arrows pointing the quiescent center (QC) (lower arrow) and the first elongated cortex cell (upper arrow). **c-d.** Primary root length and shoot fresh weight, respectively. Values are from 4 independent biological replicates each including 90-96 plantlets per condition. **e-f.** RAM length was assessed both in µm (**e**) and number of cells in the cortex cell layer (**f**). **g.** Mean CC size in the RAM, obtained by dividing the length of RAM by the number of cortex cells for each meristem. **e-g**. Data were pooled from 2 independent experiments (representing 24 individual RAM per condition). **c-g.** Black dots and whiskers represent mean values and standard deviations, respectively. Different letters correspond to statistically different groups (ANOVA type I, *p-value* < 0.05).

Additionally, upon Zn deficiency, the elongation of cortex cells was initiated closer to the tip of the root, thus resulting in an increased cortex cell size compared to control roots at the same relative position in the elongation zone (**Fig. 2a**). This difference was striking within the first 250 µm adjacent to the RAM, where the elongation was estimated to be ∼2x faster (**Fig. S2**), yielding 2.7 times longer cells at a distance of 500 µm from the QC in Zn deficiency-exposed roots (**Fig. 2a**). In contrast, mature cortex cells were ∼13% smaller upon Zn deficiency (**Fig. 2b**). Finally, cell differentiation initiated closer to the tip of the root, as the appearance of the first root hairs was closer to the QC (**Fig. 2c**).

**Figure 2.**
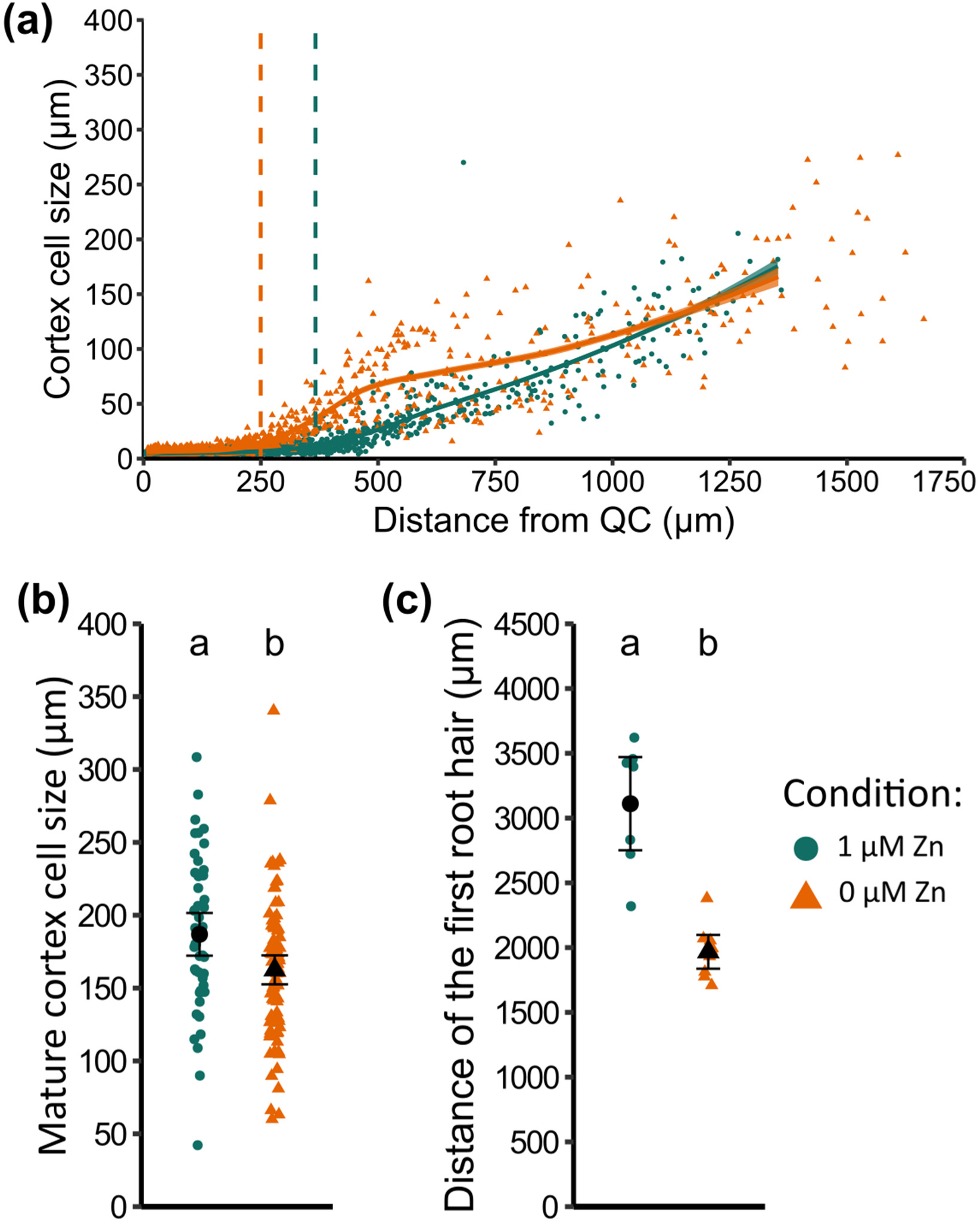
Effect of Zn deficiency on cell elongation. Plantlets were grown for two weeks on EDTA-washed agar plates in control (1 µM Zn) or Zn deficiency (0 µM Zn) conditions. **a.** Distribution of cortex cell size as a function of the distance from the quiescent center (QC). The vertical dashed lines represent the calculated mean RAM size for each condition. The trend lines and SE (standard error) areas were calculated with the “loess” function of ggplot2 R package (x∼y) with a SE confidence interval of 0.95, and span of 0.5. **b.** Mean length of mature cortex cells, measured at a distance between 5 and 6 mm from the tip of the root. **a-b**. Data from 2 independent experiments (each 12 individual primary RAM per condition). Either all the visible cortex cells (**a**) or three representative elongated cortex cells were measured between 5 and 6 mm from the QC for each root (**b**). **c**. Elongation zone (EZ) size measured as the distance between the meristem limit and the first root hair. Black dots and whiskers represent mean value and standard deviation, respectively. Different letters correspond to statistically different groups (ANOVA type I, *p-value* < 0.05).

Altogether, Zn deficiency had an impact on RAM morphology, potentially affecting three developmental processes: division, elongation and differentiation.

Observation of propidium iodide-stained roots did not indicate cell death occurring in RT exposed to Zn deficiency, as the absence of propidium iodide staining inside meristematic cells suggested that plasma membrane integrity was preserved (see Material and Methods, **Fig. 1b**). This observation, however, did not exclude compromised meristematic activity and/or quiescence upon Zn deficiency. In pWOX5:GUS lines, reporting for quiescence maintenance (Sarkar *et al*., 2007), the intensity and localization of the GUS staining were unaffected by Zn deficiency (**Fig. 3a**), indicating that both the QC identity and function were maintained upon Zn deficiency.

**Figure 3.**
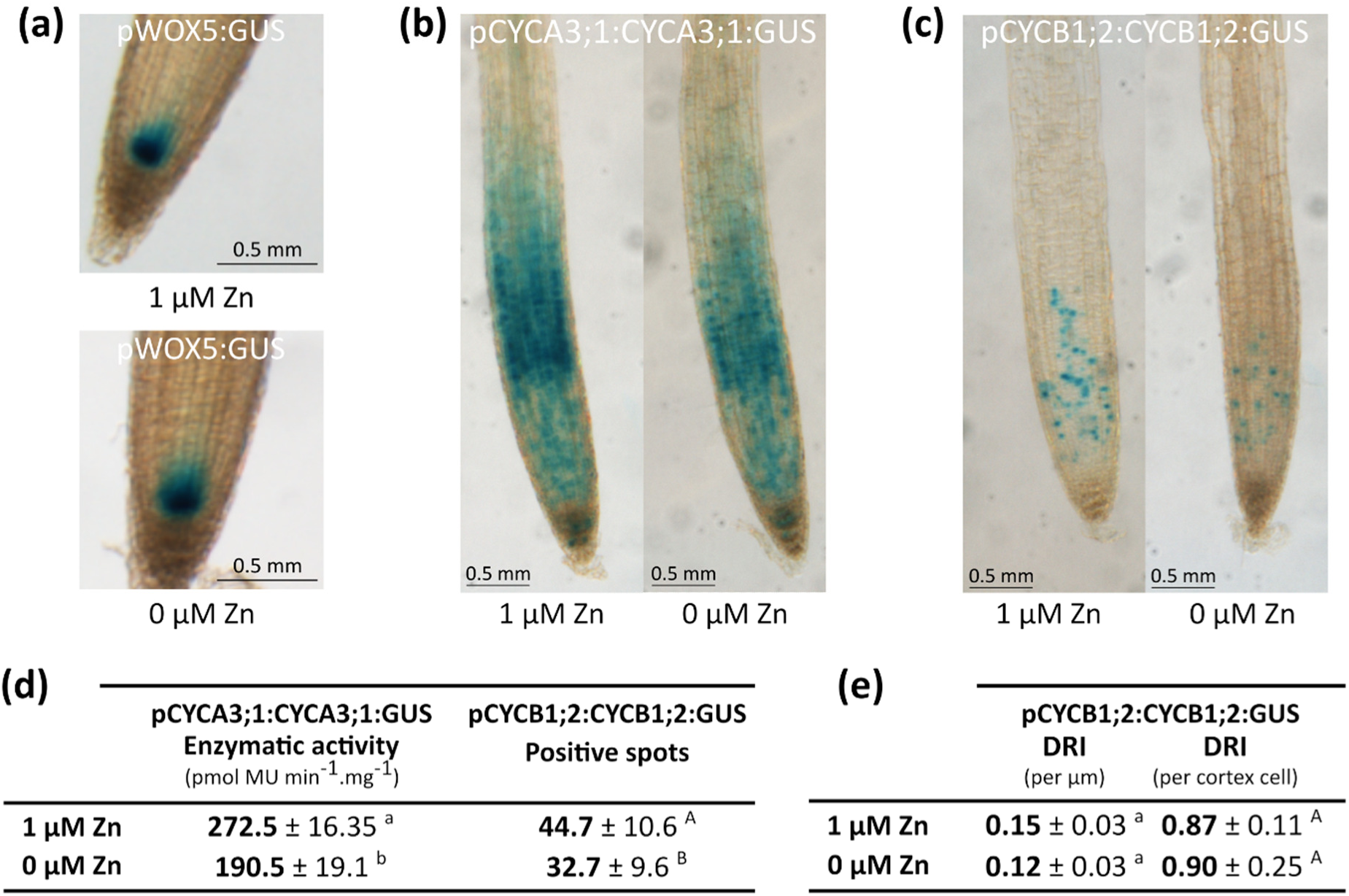
Effect of Zn deficiency on the root apical meristem (RAM) activity. Plantlets were grown for two weeks on EDTA-washed agar plates in control (1 µM Zn) or Zn deficiency (0 µM Zn) conditions. **a-c.** Representative GUS staining of pWOX5:GUS (**a**), pCYCB1;2:CYCB1;2:GUS (**b**) and pCYCA3;1:CYCA3;1:GUS (**c**) plantlets. GUS staining was performed in biological triplicates (5 plantlets each) per reporter line and conditions. **d.** Quantification of the GUS activity in the root tips of pCYCA3;1:CYCA3;1:GUS (MUG assay, in biological triplicates with samples each composed of ∼100 root tips) and pCYCB1;2:CYCB1;2:GUS (number of GUS spots; 15 plantlets per condition). **e.** Division Rate Index (DRI) for pCYCB1;2:CYCB1;2:GUS root tips (i.e. activity/RAM length), as described in Perilli & Sabatini (2010). **d-e**. Different letters correspond to significantly different groups (ANOVA type I, *p-value* < 0.05).

Next, the progression of the cell cycle was assessed by exposing pCYCA3;2:CYCA3;2:GUS (S phase reporter) and pCYCB1;2:CYCB1;2:GUS (G2/M phase reporter) lines, both expressing a fusion protein susceptible to post-translational modifications (Cools *et al*., 2010; Bulankova *et al*., 2013; Schnittger & De Veylder, 2018), to Zn deficiency (**Fig. 3b-c**). The GUS staining localization was conserved in both lines across all tissues in Zn-deficient RAM, compared to control RAM (**Fig. 3b-c**). However, quantification of the GUS staining revealed that both cell cycle markers were ∼30% less expressed in Zn deficiency-exposed roots, suggesting a global effect on cell division in the RAM (**Fig. 3d**).

Despite this observation, computing the Division Rate Index (Perilli & Sabatini, 2010) to assess the effect of Zn deficiency on the division activity of individual cells in the meristem, did not reveal any difference between Zn deficiency and control conditions (**Fig. 3e**). This suggests that individual cells underwent both S and G2/M cell cycle phases at the same rate, in both conditions. The duration of cell cycle phases was further assessed upon hydroxyurea (HU)-induced synchronization, and monitoring of re-entry in S and G2/M phases over 28 hours (**Fig. S3a-b**). In plantlets grown in control conditions, GUS staining peaked at 12-16h (**Fig. S3a**) and 20-24h (**Fig. S3b**) post HU treatment, for S and G2/M phase markers, respectively. At the temporal resolution used in this experiment, no alteration of the timing of S and G2/M phase progressions was observed in plantlets grown in Zn deficiency, although a reduced number of GUS-stained cells was observed at certain timepoints for the G2/M lines (**Fig. S3c**). Altogether, these findings suggested that the reduced RAM size mainly resulted from a reduced number of dividing cells (**Fig. 3b-d**). This was caused either by accelerated elongation rather than from reduced division rate per cell, or by a defect in cell cycle progression within the division zone. Additionally, Zn deficiency led to shorter mature cells (**Fig. 2b**), and differentiation initiating closer to the tip of the root (**Fig. 2c**).

### Zinc distribution differs in RAM and differentiated root tissues

Zn distribution in Col-0 root tissues was examined using LA-ICP-MS (Persson *et al*., 2016; Thiébaut *et al*., 2025). Transversal sections were acquired across fully differentiated roots and the primary RAM (**Fig. 4).** The ^39^K distribution was mainly used to verify the integrity of the root sections: it was homogeneous across every cell layer of the root, except for a few cortex cells in the differentiated root sections, which sometimes appeared empty in their centre (**Fig. S4**).

**Figure 4.**
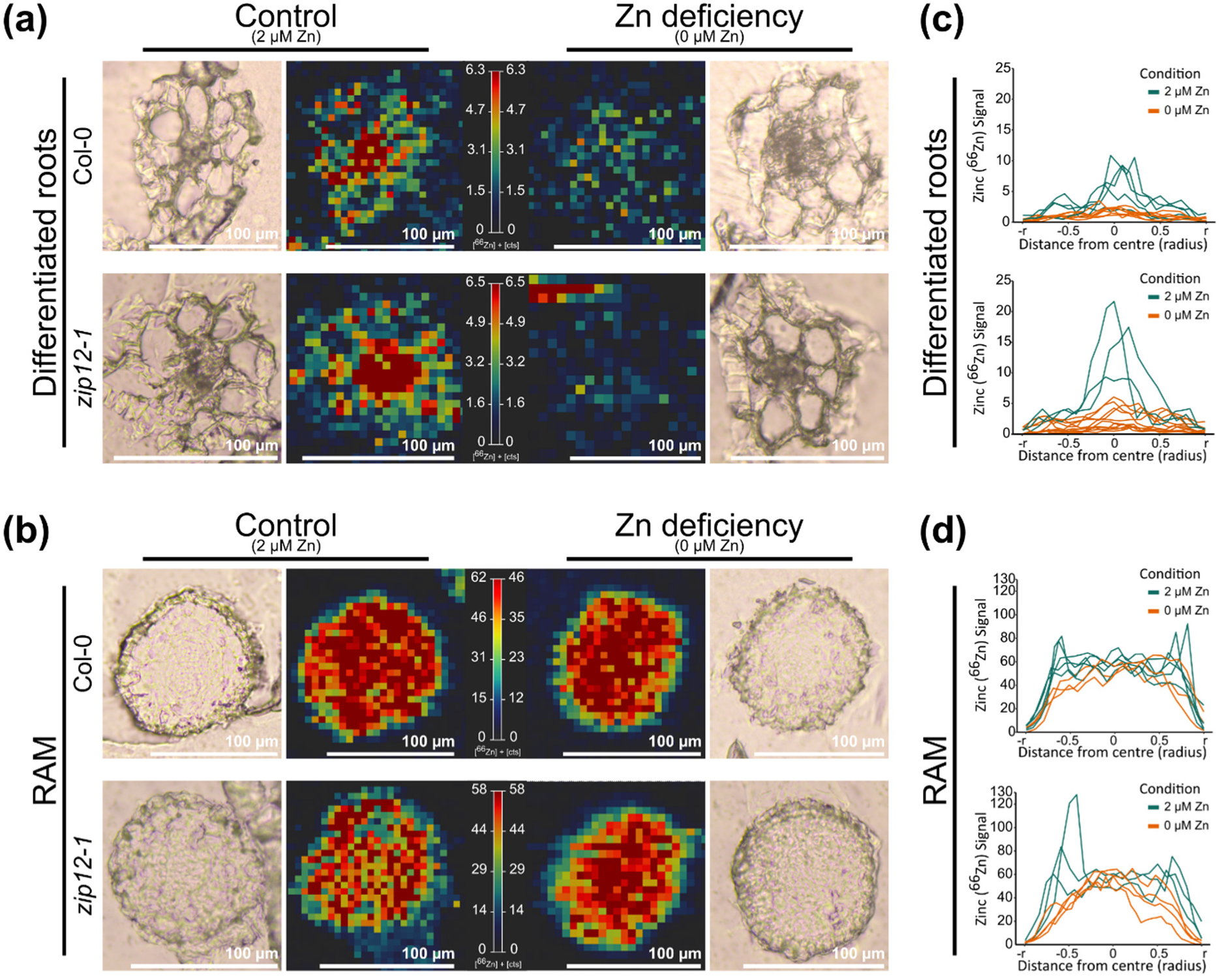
Zn distribution in the root apical meristem (RAM) and differentiated roots upon Zn deficiency in Col-0 and *zip12-1*. Col-0 and *zip12-1* plantlets were grown for two weeks on EDTA-washed agar plates in control (2 µM Zn) or Zn deficiency (0 µM Zn) conditions. Differentiated root (**a, c**) and RAM (**b, d**) transversal sections were then analysed by Laser Ablation ICP-MS. **a-b**. Representative images of the Zn localization. The Zn signal (middle) is surrounded by section optical images (left and right). Laser ablation ICP-MS was conducted on 3 independent replicates of 4 plantlets each, per condition and root part. **c-d**. Intensity graphs representing the Zn signal intensity across individual differentiated roots (**c**) or RAM (**d**) sections. Each graph contains signal from 3-6 well-dried sections, from 3 independently grown plantlets.

In control conditions, the ^66^Zn distribution peaked in the vascular cylinder of differentiated roots (**Fig. 4a**).The ^66^Zn signal was higher in the RAM than in differentiated roots, with a more homogenous distribution across all cell layers and a slight tendency for higher ^66^Zn accumulation in the epidermis [**Fig. 4b**, (Thiébaut *et al*., 2025)]. Upon Zn deficiency, ^66^Zn levels strongly decreased across differentiated roots (**Fig. 4a,c**). In contrast, ^66^Zn was maintained in the RAM, at levels similar to those of the control condition (**Fig. 4b**). Zn depletion seemed to progress centripetally across cell layers, with a slightly decreasing ^66^Zn concentration in the epidermis, but with conserved Zn level across the other tissues of the RAM (**Fig. 4b,d**). Interestingly, the ability of the RAM to maintain cellular Zn levels mirrored the maintained meristematic activity upon Zn deficiency.

### Root tip and remaining root display distinct transcriptomic responses to Zn deficiency

To capture the impact of Zn deficiency on root tip (RT) processes, including distinct Zn homeostasis, the transcriptional response of the RT to Zn deficiency relative to the remaining root (RR) system was examined by RNA-Seq analysis (Richtmann *et al*., 2025; Thiébaut *et al*., 2025). Again, Arabidopsis plantlets were germinated and grown for two weeks on control (1 µM Zn) or Zn-deprived (0 µM Zn) agar medium. At harvest, RT (∼2 mm from the tip of the root, including columella, RAM and elongation zone) of the primary roots were separated from the RR, prior to sample preparation and RNA-Seq analysis.

In agreement with previous studies using the same experimental design (Richtmann *et al*., 2025; Thiébaut *et al*., 2025), the root part (RT vs RR) explained ∼90% of the variance in gene expression, whereas the Zn deficiency treatment explained ∼7% of this variance (**Fig. 5a**). Accordingly, 2530 and 3595 differentially expressed genes [DEGs, log_2_ (fold change) < −1 of > 1; adjusted *p-value* < 0.05] were identified between RT and RR, in control and Zn deficiency conditions, respectively (**Fig. 5b-c, Table S3**). Among those, 1825 were common to both growth conditions. The DEGs between RT and RR represented a vast range of functions according to Gene Ontology (GO) enrichment analysis [*e.g.* cell wall biogenesis and modification, root hair differentiation, cell growth or nucleic acid synthesis and transcription] (**Tables S4**, **S5**).

**Figure 5.**
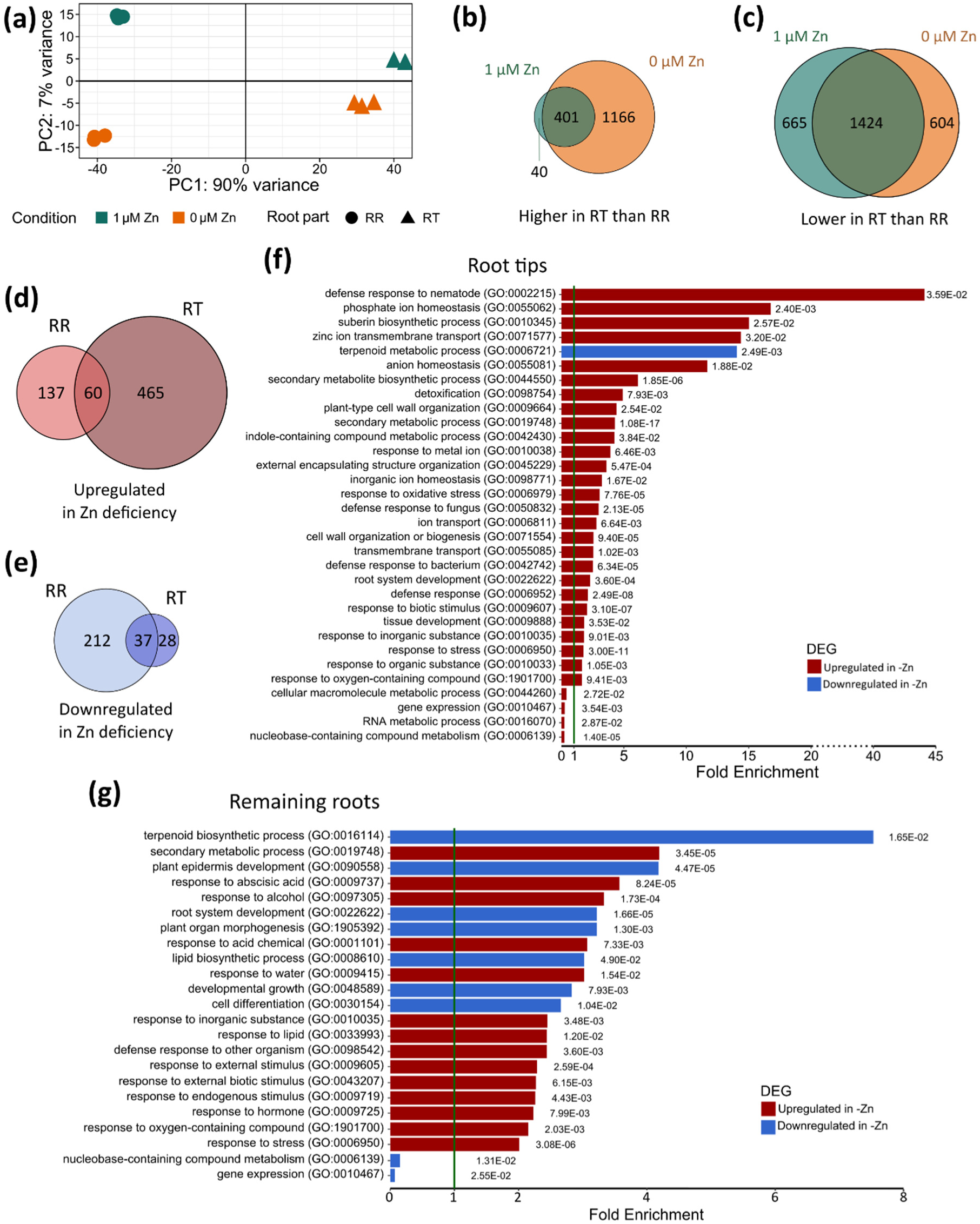
Transcriptome profiling of the root tip and the remaining root system upon Zn deficiency in Arabidopsis. Plantlets were grown in three replicates for two weeks on EDTA-washed agar plates in control (1 µM Zn) or Zn deficiency (0 µM Zn) conditions. **a.** Principal component analysis of the RNA-Seq data. **b-c.** Number of genes that are more (**b**) or less (**c**) expressed in RT than in RR samples in control (green) and Zn deficiency (orange) conditions, respectively. **d-e.** Number of genes that are up-(**d**) or down-(**e**) regulated upon Zn deficiency compared to the control condition in RT and RR samples, respectively. **b-e**. Differentially expressed genes (DEG) were identified with the following criteria: log_2_ (fold change) < −1 of > 1; adjusted *p-value* < 0.05. **f-g.** Representative selection of enriched GO terms among DEGs that are up- or down-regulated upon Zn deficiency in RT (**f**) and RR (**g**). A complete description of the GO enrichment analysis is presented in **Tables S4-S5**.

Furthermore, detailed analysis revealed distinct responses of the two root parts (RT and RR) to Zn deficiency, with only partially overlapping sets of DEGs (**Fig. 5d-e**). For instance, only 2 of the top 10 upregulated DEGs in RT upon Zn deficiency were also differentially expressed in RR (**Table 1**). Moreover, a majority of DEGs were up-regulated upon Zn deficiency in the RT (525 up vs 65 down), whereas the opposite trend was observed in the RR (249 down vs 197 up) (**Fig. 5d-e**, **Table S6**). GO enrichment analysis identified a wide diversity of up- or down-regulated processes in RT and RR upon Zn deficiency (**Fig. 5f-g**).

**Table 1.** Top differentially expressed genes (DEGs) in RT upon Zn deficiency.

| Gene | Gene name | Log<sub>2</sub> (FC) RT | *p*<sub>adj</sub> | Log<sub>2</sub> (FC) RR | *p*<sub>adj</sub> |
| --- | --- | --- | --- | --- | --- |
| *Top 10 up-regulated DEGs upon Zn deficiency in RT* |  |  |  |  |  |
| AT5G62160 | *ZIP12* | 11.76 | 1.76E-16 | 7.34 | 3.88E-76 |
| AT4G28850 | *XTH26* | 10.28 | 4.13E-11 | 1.58 | 1 |
| AT1G61080 | - | 9.60 | 4.50E-10 | 2.11 | 0.04 |
| AT3G46340 | - | 9.60 | 4.16E-10 | 0.44 | 1 |
| AT3G46370 | - | 9.55 | 1.14E-09 | 1.14 | 1 |
| AT1G51870 | - | 4 | 9.61E-09 | 1.63 | 1 |
| AT1G34047 | *DEFL208* | 9.28 | 2.24E-70 | 1.55 | 1 |
| AT3G46410 | - | 8.51 | 5.26E-06 | 3.18 | 0.29 |
| AT2G29000 | - | 8.19 | 2.55E-09 | 1.38 | 1 |
| AT5G65980 | *PILS7* | 8.19 | 3.6E-07 | 0.90 | 1 |
| *Top 10 down-regulated DEG upon Zn deficiency in RT* |  |  |  |  |  |
| AT3G60470 | - | -7.7 | 4.66E-06 | -1.43 | 1 |
| AT4G39000 | *GH9B17* | -6.54 | 3.99E-03 | -3.54 | 0.60 |
| AT3G45720 | - | -6.42 | 4.77E-12 | -6.18 | 2.50E-03 |
| AT3G12230 | *SCPL14* | -6.29 | 1.23E-14 | -4.35 | 3.97E-12 |
| AT3G28390 | *ABCB18* | -5.82 | 0.02 | -2.67 | 1 |
| AT4G09780 | - | -5.6 | 1.70E-03 | -3.15 | 2.25E-03 |
| AT3G28380 | *ABCB17* | -5.49 | 2.92E-05 | -3.95 | 1.08E-05 |
| AT1G15610 | - | -5.13 | 1.06E-05 | -3.9 | 8.71E-04 |
| AT1G78440 | *ATGA2OX1* | -4.75 | 2.20E-03 | -6.53 | 1.74E-04 |
| AT3G21670 | - | -4.68 | 2.42E-14 | 0.93 | 1 |
| *Top 10 up-regulated TF upon Zn deficiency in RT* |  |  |  |  |  |
| AT5G61470 | - | 6.58 | 7.10E-04 | 2.52 | 0.24 |
| AT3G56970 | *bHLH38* | 6.56 | 6.71E-04 | 2.54 | 2.55E-03 |
| AT2G25820 | *ESE2* | 5.91 | 3.49E-02 | 2.58 | 1 |
| AT2G44745 | *WRKY12* | 5.88 | 3.27E-02 | 5.54 | 7.89E-02 |
| AT4G28800 | - | 5.51 | 1.61E-02 | 0.66 | 1 |
| AT5G21120 | *EIL2* | 4.89 | 4.97E-02 | -0.70 | 1 |
| AT5G62380 | *NAC101* | 4.57 | 6.68E-03 | -0.17 | 1 |
| AT3G23250 | *MYB15* | 4.22 | 2.58E-14 | 1.50 | 1 |
| AT1G74650 | *MYB31* | 3.97 | 2.30E-08 | 1.19 | 1 |
| AT5G04340 | *ZAT6* | 3.94 | 5.07E-16 | 0.85 | 1 |
| *Down-regulated TF upon Zn deficiency in RT* |  |  |  |  |  |
| AT5G01900 | *WRKY62* | -4,25 | 2,08E-19 | -2,17 | 0,01 |
| AT1G65330 | *PHE1* | -3,68 | 0,02 | -4,42 | 0,37 |
| AT3G57920 | *SPL15* | -2,57 | 0,02 | -2,13 | 0,28 |
| AT2G45660 | *AGL20* | -2,19 | 2,13E-06 | -2,69 | 1,05E-11 |

Gene = Arabidopsis Gene Identifier from the Arabidopsis Genome Initiative, FC= Fold Change between Zn deficient (0 μM Zn) and control (1 μM Zn) samples, *p*<sub>adj</sub>= adjusted *p*-value (see Material and Methods): RT= Root Tips, RR=Remaining Root, TF = transcription factor.

Among the genes up-regulated by Zn deficiency in RT and RR, several over-represented GO categories were related to biotic stress responses and to interactions between organisms (**Fig. 5f-g**). Most of them were also associated with hormonal response GOs, response to hypoxia and/or to oxygen compound GOs, all belonging to general regulatory pathways, or to the response to diverse abiotic stresses including metal imbalance (Remans *et al*., 2012; Huang *et al*., 2019; Scheepers *et al*., 2020; Yaqoob *et al*., 2022; Thiébaut *et al*., 2025), suggesting common signalling of those responses.

The GO enrichment analysis further indicated that development biological processes were differentially affected by Zn deficiency in RT and RR (**Fig. 5f-g**). In RT, cell wall biogenesis and root system development were over-represented among the Zn deficiency up-regulated DEGs, while GOs associated to gene expression and RNA metabolic processes were under-represented in the same group of genes. Accordingly, among the top regulated genes in the Zn-deficient RT were found genes involved in cell wall differentiation (*XTH26*), and in the control of root development (*PILS7*, *ESE2, WRKY1*, *EIL2*) [**Table 1**, (Wang *et al*., 2007; Somssich *et al*., 2016; Wu *et al*., 2022)]. In RR, a mirrored situation was observed: root developmental processes and cell differentiation were over-represented among Zn deficiency down-regulated DEGs, while GOs associated to gene expression were under-represented in the same group of genes (**Fig. 5f-g**).

In the RT, Zn deficiency also led to the upregulation of several DEGs related to ion transport and nutrition, including enriched GOs related to phosphate (Pi) and Zn homeostasis (**Fig. 5f**). Hence, *ZIP12*, encoding a putative Zn transporter (Milner *et al*., 2013), was the top upregulated gene in RT upon Zn deficiency, and a defensin-like (DEFL208)-encoding gene which was recently linked to the impact of Zn deficiency on root growth (Kimura *et al*., 2023) was also strongly up-regulated in RT (**Table 1**). Moreover, actors involved in Fe homeostasis regulation (*bHLH38*, *WRKY12*) as well as Fe transport (*IRT2*, *VIT-like*) were found among the DEGs in the RT upon Zn deficiency (**Tables 1**, **S6**). Surprisingly, no enrichment for Zn homeostasis GOs was observed among the DEGs in the RR upon Zn deficiency (**Fig. 5f**).

Finally, top up- and down-regulated genes in RT upon Zn deficiency also included several poorly annotated and characterized genes, among them genes encoding putative kinases and transmembrane or transporter proteins (**Table 1**).

### The impact of Zn deficiency on the cell cycle and root development is reflected in the transcriptome

Root development was affected by Zn deficiency, as observed at both phenotypic (**Fig. 1-3**) and transcriptional levels with enriched development and differentiation GOs (**Fig. 5**). Root growth and development depend on cell division, but also on cell cycle exit into elongation and differentiation, often accompanied by endoreduplication (Veylder *et al*., 2011; Braidwood *et al*., 2014; Bhosale *et al*., 2018; Shen *et al*., 2025).

Although a reduced number of dividing cells was observed in the RAM upon Zn deficiency (**Fig. 3**), cell cycle processes were hardly represented among Zn deficiency-induced DEGs (**Fig. 5f-g**, **Tables 1**, **S6**). Nevertheless, hierarchical clustering of a manually curated list of cell cycle, endocycle and DNA damage response (DDR) genes (125 genes, **Table S2**, **Fig. S5a-c**) indicated (i) an overall decreased expression of endocycle and S phase genes in RR compared to RT, (ii) a decreased expression of G2/M phase genes in RT, and increased expression of DDR genes in RT and RR upon Zn deficiency, respectively (**Fig. S5**). Interestingly, root growth was negatively impacted by Zn deficiency in an *atm-1* (*ataxia telangiectasia mutated*) mutant, defective in DDR signalling (Garcia *et al*., 2003) (**Fig. S5d**).

The impact of Zn deficiency on the cell cycle was confirmed by cross-referencing Zn deficiency DEGs with a set of genes specifically expressed at the different phases of mitosis or during endoreduplication (which is linked to the onset of elongation) (**Fig. S6a**, **Table S7 Sheets 1-2**), as identified by single-cell (sc)RNA-Seq (Torii *et al*., 2020). This comparison showed an enrichment of 104 “G1-endocycle” genes specifically among the 525 DEGs up-regulated in RT upon Zn deficiency, (representing 24% of the scRNA-Seq cluster specific to this phase), mirrored by an increase of the down-regulated DEGs in RR upon Zn deficiency (**Fig. S6a,** underlined in **Table S7 Sheets 1-2**). This indicated a shift of the G1 endoreduplication gene expression from RR to RT upon Zn deficiency, which corroborated the morphological analysis marked by a shortening of the division and elongation zones (**Figs. 1**-**2**).

Finally, to further describe how Zn deficiency affects specific root cell layers, DEGs between RT and RR samples under both control and Zn deficiency conditions were compared with cell type-specific gene expression clusters from a publicly available Arabidopsis scRNA-Seq dataset. This dataset, created by Shahan *et al*. (2022), covers three developmental stages: meristematic, elongating or differentiating (**Table S7 Sheets 3-4, Fig. S6b**). The analysis showed that the elongation and differentiation gene clusters were more prominently represented in the RT at 0 µM Zn (orange curves) than at 1 µM Zn (green curves) in several cell types, specifically, in trichoblast, cortex and endodermis cells (**Fig. S6b**). These findings suggested that Zn deficiency accelerated elongation and differentiation of different RT cell layers at the molecular level. Specifically, out of the 1166 DEGs with higher expression in RT than in RR solely upon Zn deficiency (**Fig. 5b**), 444 were markers of the elongation and differentiation [(Shahan *et al*., 2022) (underlined in **Table S7 Sheets 3-4**)].

### Zn transporter genes are induced in the root tip upon severe Zn deficiency

Arabidopsis roots typically respond to Zn deficiency through the bZIP19 and bZIP23-mediated massive induction of several ZIP transporter and NAS-encoding genes (van de Mortel *et al*., 2006; Assunção *et al*., 2010; Sinclair & Krämer, 2012; Lilay *et al*., 2021; Thiébaut & Hanikenne, 2022). Several of these genes displayed increased expression in RR and in RT upon Zn deficiency, except for *ZIP10* which displayed almost no expression (**Fig. 6a**). However, only *ZIP12* passed the log_2_ (fold change) > 1 and 0.05 *p-value* thresholds in both RR and RT, whereas several additional bZIP19/bZIP23 target genes (*ZIP9*, *ZIP4*, *ZIP1*, *NAS4*) passed these thresholds solely in RT and not in RR (**Figs. 6a, S7a**). At closer inspection, these genes displayed an unusually high expression in control conditions (1 µM Zn), ranking them among the most expressed genes (**Fig. S7b**), and yet still displayed an increased expression upon Zn deficiency (0 µM Zn) in RT.

**Figure 6.**
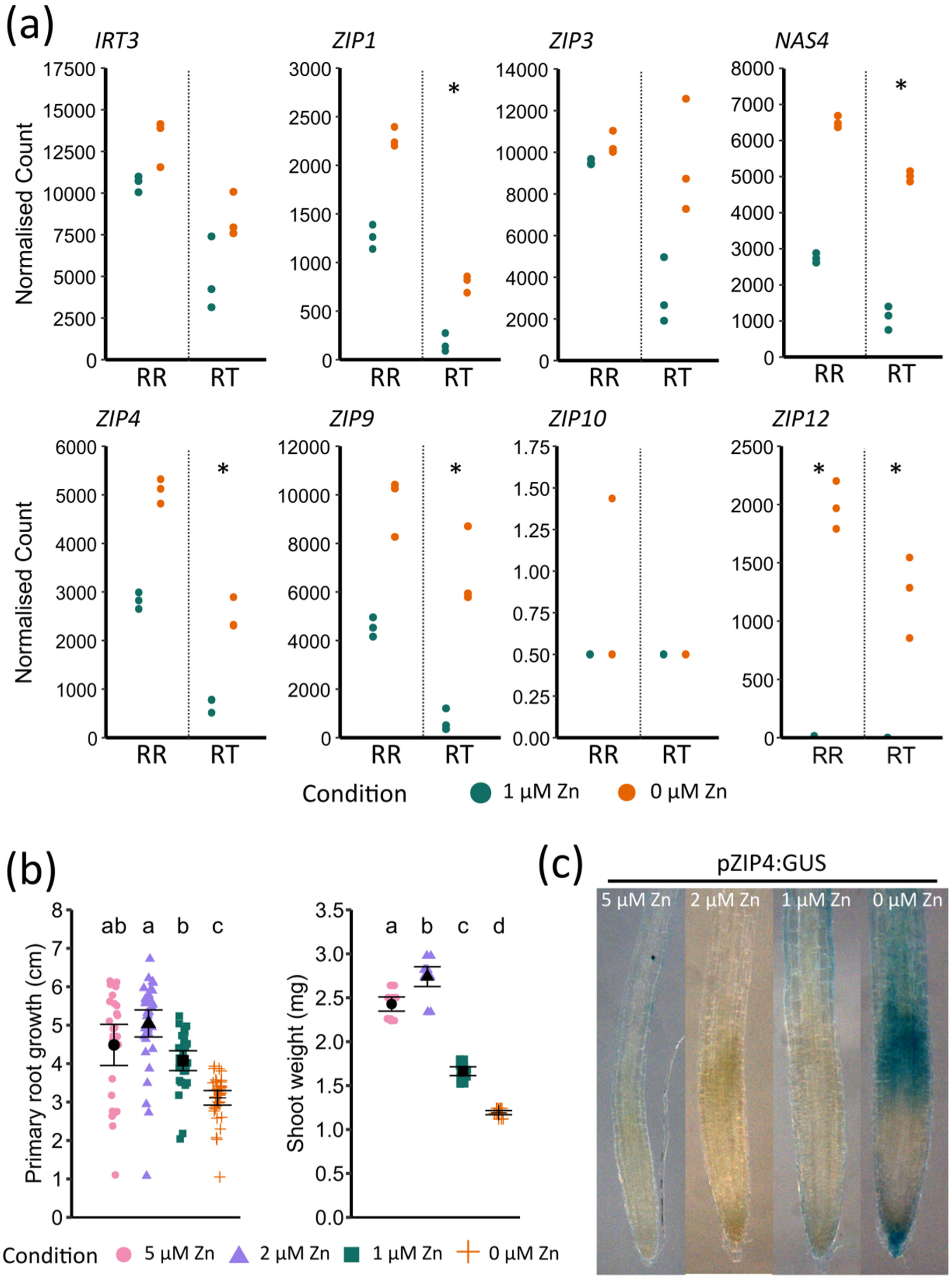
Expression of ZIP transporter genes upon Zn deficiency. Plantlets were grown for two weeks on EDTA-washed agar plates in the presence of different Zn concentrations (5, 2, 1 or 0 µM Zn, as indicated). **a**. Expression of seven ZIP transporter genes from the RNA-Seq analysis. Values are normalized read counts. Stars indicate significantly different expression between the control (1 µM Zn) and Zn deficiency (0 µM Zn) conditions (log_2_(fold change) > 1, adjusted *p-value* < 0.05). **b**. Primary root growth and shoot fresh weight of Col-0 plantlets. Values are from 3 biological replicates, each including 12 plantlets per condition. Black dots and whiskers represent mean values and standard deviations, respectively. Different letters correspond to statistically different groups (ANOVA type II, with Tukey test correction, *p-value* < 0.05). **c.** GUS staining in primary root apex of pZIP4:GUS plantlets. Pictures are representative of observations made in 10 plantlets.

Growing Col-0 plantlets across an increasing Zn concentration range (0-5 µM Zn) showed that root and shoot growth was reduced at 1 µM Zn compared to 2 and 5 µM Zn, with even greater inhibition at 0 µM Zn (**Fig. 6b**). At 2 µM Zn, several *ZIP* genes were significantly less expressed in roots, as expected for Zn sufficient conditions, compared to 0 µM Zn (**Fig. S7c**). Similarly, when growing a Zn deficiency reporter line [pZIP4:GUS, (Assunção *et al*., 2010)], GUS staining was observed in roots of plantlets grown at 1 µM Zn but not at 2 or 5 µM Zn (**Fig. 6c**). A more intense GUS staining was however observed at 0 µM Zn, particularly in the vicinity of the RAM, compared to the 1 µM Zn condition.

Agar is a source of metal nutrients (Gruber *et al*., 2013) and to achieve effective Zn deficiency in agar-based media, Zn traces must be removed, typically by washing the agar with EDTA (Sinclair & Krämer, 2012). Our observations (**Figs. 6, S7**) indicated that the plantlets grown under our initial control conditions (Hoagland media with 1 µM Zn in EDTA-washed agar) already displayed a Zn deficiency response, that was further induced at 0 µM Zn, particularly in the RT. Therefore, the 1 µM Zn condition may rather be referred to as Zn limitation (Moseley *et al*., 2002; Esteves *et al*., 2023), and 0 µM Zn as Zn deficiency. Practically, it suggested that the root morphological features and gene expression patterns reported here at 0 µM Zn compared to 1 µM Zn were those associated with a severe Zn deficiency. In follow-up experiments, 2 µM Zn was used as control, Zn replete, condition.

### ZIP12 plays a key role in the Zn deficiency response

Being very poorly expressed in control conditions, *ZIP12* was the most strongly up-regulated gene in RT (log_2_ FC = 11.8) and the 4^th^ in RR (log_2_ FC = 7.3) upon Zn deficiency (**Tables 1**, **S6**, **Fig. 6a**). *ZIP12* encodes a putative Zn transporter, as determined by yeast complementation assays (Milner *et al*., 2013) and is a target of the bZIP19 and bZIP23 TFs (Assunção *et al*., 2010). To further examine the function of *ZIP12* in Zn homeostasis, two *zip12* T-DNA insertion mutant lines, *zip12-1* (SALK_137184) and *zip12-2* (SALK_118705) (Inaba *et al*., 2015), were phenotyped in Zn deficient conditions (**Figs. 7**, **S1a**). A greater inhibition of the primary root growth upon Zn deficiency was observed in *zip12-1* compared to Col-0 and *zip12-2* (**Figs. 7a, S1b-c**). These phenotypes correlated with the *ZIP12* expression level in the roots of the 3 genotypes upon Zn deficiency, with a strong induction in Col-0 and *zip12-2*, and no expression in *zip12-1* (**Fig. 7b, d**).This suggested that the T-DNA insertion in the *ZIP12* promoter region upstream of the two ZDREs in the *zip12-2* mutant (**Fig. S1a**) did not knock-out its transcription upon Zn deficiency. Therefore, phenotyping was pursued solely with the *zip12-1* mutant. No difference in RAM size was observed between Col-0 and *zip12-1,* but the RAM cortex cells were significantly longer in *zip12-1* compared to Col-0 upon Zn deficiency (**Fig. 7c**).

**Figure 7.**
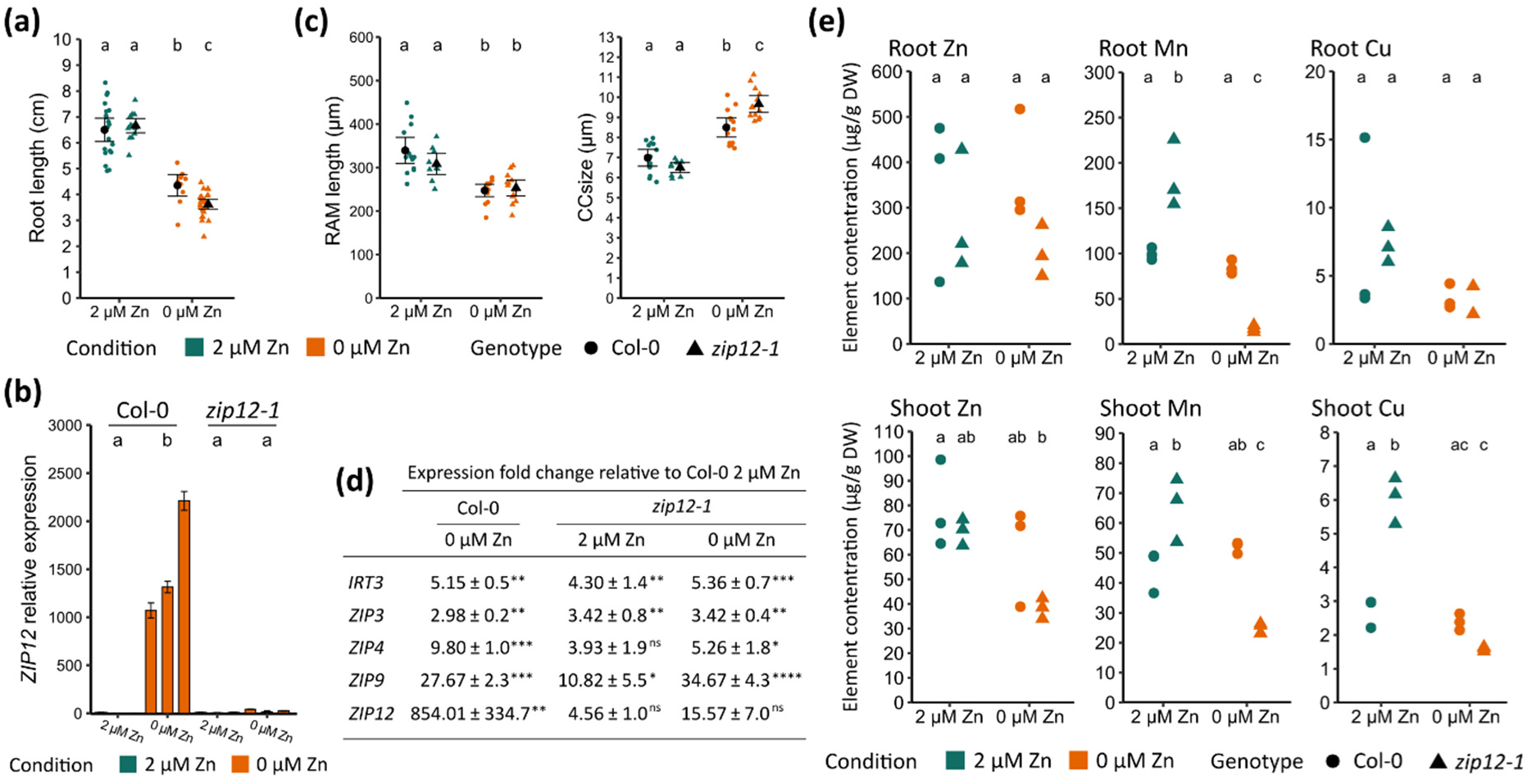
Phenotypes of the *zip12-1* mutant upon Zn deficiency. Plantlets of Col-0 and *zip12-1* were grown for two weeks on EDTA-washed agar plates in control (2 µM Zn) or Zn deficiency (0 µM Zn) conditions. **a.** Primary root length. **b**. *ZIP12* relative expression in Col-0 and *zip12-1* plantlets, as determined by RT-qPCR. Bar values and errors are means +/− SD from 3 technical replicates, normalized on mean expression in Col-0 roots (2 µM Zn). The values for three biological replicates (each including 180 plantlets) are presented. **c.** RAM length and cortex cell size in the RAM of Col-0 and *zip12-1* plantlets. **d.** Summary of the expression of ZIP transporter genes by RT-qPCR, expressed in fold change relative to their expression in roots of Col-0 seedling grown in the presence of 2 µM Zn. Values are from three biological replicates (each including 180 plantlets). Stars correspond to statistically different expression compared to Col-0 roots at 2µM Zn (ANOVA type I, *p-value* < 0.05). **e.** Root and shoot ionome profiling in Col-0 and *zip12-1.* Values are from 3 biological replicates (each including **160-180** plantlets). Different letters represent statistically different groups (ANOVA type II, with Tukey test correction, *p-value* < 0.05).

In *zip12-1*, the absence of *ZIP12* transcripts was compensated by higher expression of other *ZIP* genes (*IRT3*, *ZIP3*, *ZIP4*, *ZIP9*) in control conditions (2 µM Zn) compared to Col-0, reaching levels close to their peak expression in Zn deficiency (**Fig. 7d**). Zn accumulation in whole roots and shoot of *zip12-1* varied only marginally from Col-0 both in control and Zn-deficient conditions (**Fig. 7e**). In contrast, under control conditions, *zip12-1* accumulated 1.5 and 2.2 times more Mn and Cu in shoots, and 1.85 times more Mn in roots compared to Col-0 (**Fig. 7e**). Upon Zn deficiency, Cu levels in *zip12-1* shoots decreased to Col-0 concentrations, while Mn accumulation in *zip12-1* roots and shoots was strongly reduced, by 10.78x and 2.60x, respectively, and was below Col-0 concentrations. The shoot and root concentrations of all other elements measured (Ca, Fe, K, P, Mg, Na) remained unaffected by the genotype or the treatment, except for Mg in roots, which was reduced by Zn deficiency in both Col-0 and *zip12-1* (**Fig. S8**).

The ^66^Zn distribution in *zip12-1* root tissue sections, examined using LA-ICP-MS, revealed a similar pattern to that of Col-0: upon Zn deficiency, ^66^Zn levels were strongly reduced in differentiated roots, particularly in the vascular cylinder, whereas they were maintained in the RAM (**Fig. 4**). However, two noticeable differences in the ^66^Zn distribution were observed between the *zip12-1* mutant and Col-0. First, in control conditions, *zip12-1* displayed higher ^66^Zn levels in the vascular cylinder than Col-0. Second, the centripetal decrease in Zn level from the epidermis towards the root centre observed in Col-0 was more marked in the RAM of *zip12-1* (**Fig. 4**).

## Discussion

Root growth and architecture are strongly influenced by nutrient availability, as are the RAM morphology and activity. Low nutrient availability can ultimately result in total exhaustion of quiescence and meristematic activity in the RAM (Ward *et al*., 2008; Jain *et al*., 2009; Lequeux *et al*., 2010; Gruber *et al*., 2013; Bouain *et al*., 2019). Here, we showed that severe Zn deficiency resulted in strongly impaired primary root growth and altered RAM morphology, with (i) shortened division zone, while quiescence was maintained and individual cells retained meristematic activity, (ii) shortened elongation zone made of longer cells, and (iii) accelerated differentiation resulting in shorter mature cells (**Fig. 8**). Zn was homogenously distributed in the RAM, and this distribution was maintained upon severe Zn deficiency, in contrast to RR. Finally, our data indicated an important role of the ZIP12 transporter in the response to Zn deficiency (**Fig. 8**). Altogether, our results suggested that, despite a reduction of its size and of overall root length, the RAM was actively protected from becoming Zn deficient, possibly through prioritised Zn allocation and/or retention.

**Figure 8.**
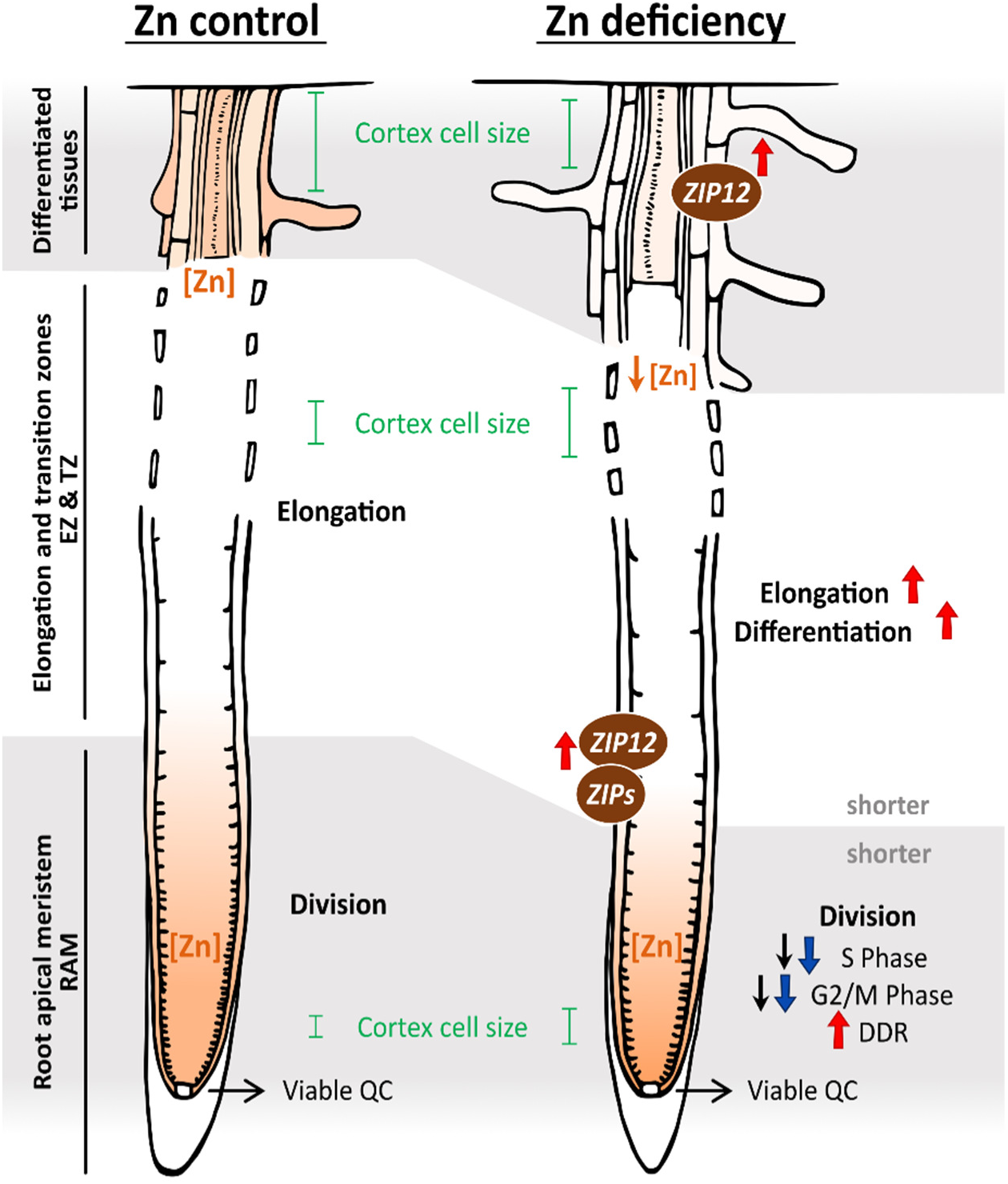
Summary of the response of the RT to Zn deficiency. Under zinc (Zn) deficiency conditions, root apical meristem (RAM) size was reduced, meristematic cortex cells were fewer and longer (Fig. 1) with their elongation starting closer to the tip of the root resulting in longer cells in the elongation zone, and differentiation occurring closer to the root apex (Fig. 2). Viability and activity of the quiescent centre (QC) and the RAM were maintained, however with fewer individual dividing cells and accelerated elongation (black arrows from GUS activity, red and blues arrows from transcriptomic data, Figs. 3, 5, Fig**. S5**), and with an increased response to DNA damage (DDR) (**Fig. S5**). Zn concentration (orange staining) strongly decreased in differentiated roots, while it was maintained in the RAM, with a slight depletion starting from the external cell layers, especially in the epidermis (Fig. 4). Finally, several *ZIP* genes were more expressed in the Zn deficiency roots, especially in the RT, with *ZIP12* being the most upregulated gene in the RT (Fig. 6, **Table 1**). RAM: Root apical meristem. EZ: Elongation zone. DZ: Differentiation zone.

### Zn deficiency alters RAM function and root zonation through multiple, interconnected mechanisms

In Arabidopsis, RAM size reduction is a common response to both nutrient deficiency and toxicity (Sánchez-Calderón *et al*., 2005; Wei *et al*., 2021; van Dijk *et al*., 2022; Thiébaut *et al*., 2025). Although RAM size, overall cell number and meristematic activity were reduced (**Figs. 1-3, 8**, **S2**), quiescence and division at the single cell level persisted, with no sign of cell death (**Figs. 1, 3, S3**). This was consistent with the observation that Zn levels in the RAM remained relatively stable under Zn deficiency (**Figs. 4, 8**). At the transcriptomic level, Zn deficiency had limited impact on core cell cycle gene expression (**Fig. S5a-c**). However, regulation of the cell cycle, for instance upon stress, is mainly a post-transcriptional process (Marrocco *et al*., 2010; Genschik *et al*., 2014; Hu *et al*., 2016; Velappan *et al*., 2017), which may account for the observed reduction in meristem activity.

Several processes are known to contribute to RAM size in stress conditions, including for instance DDR, cytoskeletal dynamics and hormonal signalling (Lipka & Müller, 2014; van Dijk *et al*., 2022; Shi *et al*., 2023). Several of these processes were impacted by Zn deficiency, as outlined below.

First, DDR pathways are known to mediate cell cycle arrest and RAM size reduction in response to stresses such as Pi deficiency or aluminium toxicity (Sánchez-Calderón *et al*., 2005; Wei *et al*., 2021). In this study, Zn deficiency did not induce cell death but did increase the expression of DDR-related genes, both in the RT and RR (**Fig. S5a-c**). Notably, an *atm* mutant, lacking a key DDR kinase, exhibited heightened sensitivity to Zn deficiency, with reduced root growth (**Fig. S5d**). The ATM kinase mediates S and G2 phases arrest and DNA repair, enabling genome protection under stress (Kurz & Lees-Miller, 2004; Nisa *et al*., 2019). This suggested that ATM may play a role in maintaining root growth under Zn deficiency conditions by enabling a protective DDR and RAM size control.

Second, Zn deficiency increased cell size within the RAM, which is a rather uncommon phenotype under nutrient stress. In Arabidopsis, metal stress-related cell expansion in the RAM was previously described upon Ni excess (Lešková *et al*., 2020) and in the Cu-sensitive *copper modified resistance1/patronus1* (*cmr1/pans1*) mutant (Juraniec *et al*., 2016). In both cases, the large cell phenotypes were linked to altered microtubule dynamics. Since Zn promotes microtubule polymerization (Hesketh, 1982; Oteiza *et al*., 1990), Zn deficiency may alter microtubule homeostasis in the RT, leading to cell expansion. This hypothesis merits further investigation. Alternatively, endoreduplication is known to contribute to cell expansion via cell wall remodelling (Takahashi, 2013; Bhosale *et al*., 2018) The higher expression of endoreduplication genes observed in the RT upon Zn deficiency (**Fig. S6a, Table S7 Sheets 1-2**) may possibly drive cell expansion in our experimental conditions.

Third, root zonation, *i.e.* the delineation of the division, elongation and differentiation zones in the RT, is tightly controlled. Here, using the Shahan *et al*. (2022) scRNA-Seq dataset, which provides cell type-specific gene expression clusters across different developmental stages, we showed that under Zn deficiency conditions, markers of elongation and differentiation were more prevalent in RT compared to Zn control conditions. This suggested an acceleration of elongation and differentiation (**Fig. S6b, Table S7 Sheets 2-4**). These processes are tightly regulated by hormonal signals (Tsukagoshi *et al*., 2010; Takahashi, 2013; Bhosale *et al*., 2018; Zluhan-Martínez *et al*., 2021). In particular, opposite auxin and cytokinin (CK) gradients are known to regulate the exit of cell cycle and thus the transition from division to elongation and differentiation zones (Chapman & Estelle, 2009; Tsukagoshi *et al*., 2010; Kong *et al*., 2018; Bhosale *et al*., 2018). Zn deficiency generally decreases auxin levels in dicot roots (Thiébaut & Hanikenne, 2022; Lilay *et al*., 2024), and genes involved in auxin signalling have been linked to Zn homeostasis in maize (Xu *et al*., 2022; Wang *et al*., 2023). In parallel, CK are also associated to the control of Zn homeostasis (Gao *et al*., 2019). In our dataset, auxin and CK signalling genes showed only moderate changes in expression, with the notable exception of the downregulation of *ARR15* (log_2_(FC) −3.52, **Table S6**), a repressor of CK signalling (Kiba *et al*., 2003). Since CK is known to promote elongation-related processes such as endocycling and cell-wall synthesis, to the detriment of cell cycle activity, meristematic cell number and RAM size (Takahashi *et al*., 2013; Takahashi & Umeda, 2014; Liu *et al*., 2022), the repression of *ARR15* upon Zn deficiency may enhance these CK-mediated processes and thereby potentially contribute to RAM shortening. In addition, ethylene is a key regulator of root development, particularly of root hair formation (Zluhan-Martínez *et al*., 2021; Martin *et al*., 2022). In this study, we observed that upon Zn deficiency, root hairs initiated closer to the RAM (**Fig. 1, 2, 8**), which is a hallmark of altered root zonation. Moreover, two ethylene-responsive transcription factors, *ESE2* and *EIL2*, were among the most strongly up-regulated TFs (log_2_ FC 5.91 and 4.89, respectively; **Table 1**). These findings support the hypothesis of a role for ethylene signalling in the Zn deficiency response, which is consistent with its involvement in other nutrient stresses (Barberon *et al*., 2016; Gutiérrez-Alanís *et al*., 2017; García *et al*., 2021)

Finally, the *DEFL202, 203, 207,* and *208* genes were strongly upregulated in the RT upon Zn deficiency (**Table 1, Table S6**). Defensin-like (DEFL) peptides, which are small cysteine-rich peptides known for their role in various biotic and abiotic stress responses (Stotz *et al*., 2009; Wu *et al*., 2016; Domingo *et al*., 2024), are additional candidates for a potential role in RAM size regulation. DEFL-encoding genes were described as part of the Zn deficiency response, possibly via the action of bZIP19 (Inaba *et al*., 2015; Kimura *et al*., 2023). Notably, a *defl202/203* double mutant exhibits longer RAM and primary root in Zn-sufficient conditions. Although the underlying mechanism of action remains unclear, DEFL peptides have been proposed to limit the RAM size as a strategy to conserve Zn resources in the RAM under low Zn availability. However, the *bzip19/23* double mutant, even though deficient in *DEFL* induction, shows shorter roots under Zn deficiency (Lilay *et al*., 2019), possibly due to impaired Zn allocation and early RAM exhaustion.

### Zn concentration is maintained in the RAM during Zn deficiency

In differentiated roots of Arabidopsis plants grown in control conditions (2 µM Zn), Zn was mostly concentrated in the vascular cylinder (**Fig. 4a, c**). This is consistent with the fact that xylem loading is key for Zn translocation to the shoot (Hussain *et al*., 2004; Sinclair *et al*., 2007) and is considered as a bottleneck for Zn translocation to the shoot (Claus *et al*., 2013; Kisko *et al*., 2018). In contrast, Zn was homogeneously distributed in the RAM, accumulating at concentration approximately ∼10-fold higher than in differentiated roots (**Fig. 4b, d**, (Ochoa Tufiño *et al*., 2025; Thiébaut *et al*., 2025), likely reflecting the primary role of Zn in active cell division (Falchuk *et al*., 1975; Francis *et al*., 1995). Despite the low expression of genes encoding ZIP transporters, considered as the primary Zn uptake system in Arabidopsis roots, in Zn sufficient condition [**Fig. S7**, (van de Mortel *et al*., 2006; Ochoa Tufiño *et al*., 2025)], the Zn enrichment in the RT suggested that Zn may be more efficiently taken up, retained in and/or actively translocated to the meristem. The higher Zn levels in future epidermal cells in the RAM further support a role for Zn uptake in shaping the Zn distribution pattern in the meristematic zone (**Figs. 4, 8**). Recently, *IRT3* and *YSL3* were shown to be expressed upon Zn deficiency in the division zone, whereas *ZIP3* and *ZIP5*, involved in Zn uptake, were expressed in epidermis cells of the elongation zone (Ochoa Tufiño *et al*., 2025).

Upon Zn deficiency, Zn levels dramatically decreased in differentiated roots (**Fig. 4a, c**). The gradual upregulation of *ZIP* genes when roots are exposed to Zn sufficiency (2 µM Zn), Zn limitation (1 µM Zn) or severe Zn deficiency (0 µM) (**Figs. 6, S7**) appeared to be insufficient to maintain Zn homeostasis when Zn supply was severely low. Such a gradual and fine-tuned response to Zn deficiency may explain the fact that Zn deficiency has been reported to have either a negative (Talukdar & Aarts, 2008; Mager *et al*., 2018; Kimura *et al*., 2023), a positive (Sinclair & Krämer, 2012; Lilay *et al*., 2019), or no visible (Assunção *et al*., 2010; Gruber *et al*., 2013) effect on primary root growth in Arabidopsis. These apparent inconsistencies among previous reports are likely linked to how much Zn traces remained in the growth medium and thus to the severity of Zn deficiency applied in the different experimental systems. The different growth media included the use of different agar powders (Jain *et al*., 2009; Gruber *et al*., 2013), either washed with EDTA, or not (Sinclair & Krämer, 2012; Charlier *et al*., 2015). Here, we observed that a control growth medium made of Hoagland with 1 µM Zn and EDTA-washed agar (i.e. with reduced Zn contamination) was not a Zn sufficient condition, resulting in Zn limitation, *i.e.* an intermediate state where root growth was only moderately affected, but where the bZIP19/bZIP23-dependent molecular response to Zn deficiency was activated already. Such a gradual response was previously reported in the alga Chlamydomonas, where the response to Fe deficiency progressed from an induction of uptake to a massive metabolic reorganization as the Fe supply was gradually decreased (Moseley *et al*., 2002; Urzica *et al*., 2012).

In contrast to RR, Zn levels were essentially maintained in the RAM in Zn deficiency conditions (0 µM Zn), as shown both by Zn imaging in the RAM (**Figs. 4b, d, 8**) and by the limited upregulation of *ZIP* genes in Zn-limiting conditions (1 µM Zn) (**Fig. 6a**). Such prioritized Zn allocation to the RT possibly preserved the meristematic activity. Several hypotheses can be formulated to interpret this observation. First, strong upregulation of several *ZIP* genes in the RT upon Zn deficiency [**Figs. 6**, **S7**, (Ochoa Tufiño *et al*., 2025)] could contribute to stronger Zn uptake and Zn distribution between cell layers. For instance, the strong increase of *pZIP4:GUS* expression in the vicinity of the RAM was striking. Despite this strong induction of Zn transport, Zn depletion was visible in the most external cell layer of the RAM, suggesting that the Zn homeostatic mechanisms were also starting to be overcome. Second, the densely organized meristem, composed of smaller cells with a reduced vacuolar lumen and thus more concentrated enzymes, molecules and cell-walls than differentiated roots, may act as a Zn sink, contributing to higher Zn concentrations in the RAM. Lastly, as observed for Pi homeostasis (Kanno *et al*., 2016), the entire root system could contribute to Zn uptake, which would then be translocated to the RAM.

Finally, in two separate studies, we observed reduced Zn and cadmium (Cd) accumulation in the Arabidopsis RAM compared to differentiated tissues upon Zn excess (Thiébaut *et al*., 2025) or Cd exposure (Richtmann *et al*., 2025), respectively. Combined, these observations indicate that Zn homeostasis in the RAM is maintained over a wider range of external Zn supply than in differentiated tissues, prioritizing Zn allocation to the RAM upon Zn deficiency or protection of the RAM upon Zn excess (or Cd exposure). This mobilizes a specific response of Zn homeostasis genes (including *ZIP12* upon Zn deficiency, see below), but also of the homeostasis of other nutrients, such as Fe or Pi, of the specialized metabolism, of cell wall modification processes and/or of developmental processes (this study; Richtmann *et al*., 2025; Thiébaut *et al*., 2025).

### The interaction between Zn and Pi homeostasis also takes place in the RT

The crosstalk between Zn and Pi homeostasis is well documented. In Arabidopsis, low Pi supply increases shoot Zn accumulation, whereas Zn deficiency induces Pi uptake (Khan *et al*., 2014; Kisko *et al*., 2018). In line with this interplay, we observed that Zn deficiency induced the expression of Pi-related genes in the RT. These include two genes encoding Pi transporters (*PHO1;H1* and *PHT3;2*) and three members of the Pi starvation-induced glycerol-3-phosphate permease gene family (*G3Pp1-3*) (**Table S6**). Additionally, *ZAT6*, a gene previously shown to be up-regulated during Pi starvation (Devaiah *et al*., 2007), was also induced in RT upon Zn deficiency (**Table 1**). The ZAT6 TF controls root development and multiple stress responses, including response to Cd exposure, acting on the redox status (Chen *et al*., 2016; Tang & Luo, 2018). Interestingly, *PHT1;1*, a gene encoding a major contributor in Pi uptake upon Zn deficiency (Ayadi *et al*., 2015; Rai *et al*., 2015), was not identified among DEGs in the RT (**Table S6**). Finally, *PILS7* (*PIN-LIKES7*), encoding a putative auxin transporter, was strongly upregulated in RT upon Zn deficiency (Log_2_ FC 8.2, **Table 1**). PILS7 has previously been shown to control root growth in response to Pi supply, and other stress conditions (Yi *et al*., 2021) suggesting it may also contribute to the Zn deficiency response.

### ZIP12 acts a last resort ZIP transporter upon Zn deficiency

The induction of *ZIP* gene expression is a hallmark of the Zn deficiency response in plants (Talke *et al*., 2006; van de Mortel *et al*., 2006; Assunção *et al*., 2010; Dong *et al*., 2018; Amini *et al*., 2022; Thiébaut & Hanikenne, 2022). In this study, several *ZIP* genes were strongly up-regulated in the RT upon Zn deficiency (**Figs. 6a**, **S7**). Among them, *ZIP12* was the most up-regulated gene in the RT. The ZIP12 transporter is a putative Zn transporter, as it was shown to complement a Zn uptake-impaired yeast mutant, but did not appear to transport other metals such as Mn or Fe (Milner *et al*., 2013). The *ZIP12* promoter contains Zn deficiency response elements (ZDRE), placing the gene under the control of the F-group bZIP TFs, and more specifically of bZIP23 (Assunção *et al*., 2010; Jain *et al*., 2013; Inaba *et al*., 2015; Lilay *et al*., 2019). The activity of ZIP12 upon Zn deficiency was recently shown to be dependent on phosphorylation by calcium-dependent CBL-CIPK (calcineurin B-like-CBL-interacting protein kinases) signaling (Fang *et al*., 2025).

Unlike most *ZIP* genes, *ZIP12* showed minimal expression in differentiated roots and RT under Zn-limiting conditions (1µM Zn) but was strongly induced upon severe Zn deficiency (0 µM Zn) (**Fig. 6a**), suggesting that it may play a specific role in these conditions. Two *zip12* mutants, *zip12-1* and *zip12-2*, were partially characterized by Inaba *et al*. (2015), both displaying reduced root growth upon Zn deficiency, despite unaltered root Zn levels (Inaba *et al*., 2015). In our study, only *zip12-1* showed impaired root growth upon Zn deficiency, while *zip12-2* retained strong *ZIP12* expression upon Zn deficiency, consistent with a T-DNA insertion upstream of the 2 ZDREs in the *ZIP12* promoter (**Fig. S1**). In addition, *zip12-1* displayed (i) altered RAM morphology, (ii) constitutive induction of several Zn-responsive *ZIP* genes, (iii) reduced shoot Zn accumulation, (iv) strongly altered Mn and Cu accumulation, (v) elevated Zn level in the vascular cylinder of differentiated roots in control conditions, and (vi) mostly unchanged Zn level in the RAM upon Zn deficiency (**Fig. 7**, **Fig. 4**). Altogether, these findings indicated that *ZIP12* plays an important role in Zn spatialization in the root. We suggest that the strong, constitutive, upregulation of several *ZIP* genes *zip12-1* was responsible for the complex phenotype of the mutant, affecting multiple metals and cell layers. It was also possibly responsible for the marginal impact of the *zip12-1* mutation on Zn levels in plantlets (this study; Inaba *et al*., 2015).

According to a recent study by Ochoa Tufiño *et*), *ZIP12* is predominantly expressed in the epidermal cell layer, in the zone connecting the root and the shoot. We showed here that *ZIP12* is also strongly induced by Zn deficiency in the RT (**Fig. 6a**). Moreover, ZIP12 is presumably located to the plasma membrane (Fang *et al*., 2025). In the *zip12-1* mutant, the concomitant *ZIP12* loss-of-function and up-regulation of *IRT3*, *ZIP4* and *ZIP9* (**Fig. 7d**), which was shown to be involved in Zn radial transport across the root (Lee *et al*., 2021), may explain both Zn depletion in the epidermis and increased Zn accumulation in the vascular cylinder of differentiated roots (**Fig. 4a, c**). Moreover, the increased Mn accumulation in roots and shoot of the *zip12-1* mutant in control conditions (**Fig. 7e**) may be explained by the upregulation of *ZIP9* (FC ∼11 compared to Col-0), encoding a putative Mn transporter (Milner *et al*., 2013). Why Mn levels were drastically down in *zip12-1* upon Zn deficiency remains to be determined. Increased Mn content upon Zn deficiency in Arabidopsis has been observed previously and was attributed to the regulatory function of the bZIP19 and bZIP23 TFs, potentially via the upregulation of ZIP transporter gene expression (Lilay *et al*., 2019). In this context, it is interesting to note that neither *ZIP12* nor *ZIP9* are known to be upregulated by Mn deficiency in Arabidopsis (Rodríguez-Celma *et al*., 2016). Finally, the constitutive upregulation of several *ZIP* genes in *zip12-1*, all with specific cell-type expression patterns (Ochoa Tufiño *et al*., 2025) and all targets of bZIP19 and bZIP23 (Assunção *et al*., 2010), suggested that these two TFs may have specific functions, sensing differential Zn status among cell types.

For a long time, knowledge regarding the metal specificity, subcellular localization and overall function of individual ZIP transporters in Arabidopsis has been lagging behind (Ricachenevsky *et al*., 2015). This mainly stems from the fact that, with a few exceptions (Milner *et al*., 2013; Inaba *et al*., 2015), single *zip* mutants typically fail to display strong phenotypes. Recently, the use of multiple mutants has started unveiling the contribution of ZIPs to the Zn uptake pathways in Arabidopsis roots (Lee *et al*., 2021; Ochoa Tufiño *et al*., 2025), highlighting the functional redundancy among ZIP transporters. Hence, a quadruple *irt3*, *zip4*, *zip6* and *zip9* mutant is defective in Zn radial transport across the roots, and in consequence displays strongly altered Zn homeostasis and seed development (Lee *et al*., 2021). A double *zip3*/*zip5* mutant is defective in Zn uptake, while Zn levels are maintained in the single *zip3* or *zip5* mutants (Ochoa Tufiño *et al*., 2025). Strikingly, despite its very low expression under Zn-replete and Zn--limiting conditions, but strong induction upon severe Zn deficiency, *ZIP12* emerges as a key actor in the Zn deficiency response. In particular, the fact that a single *zip12* mutant displayed significant alteration in both growth and metal homeostasis points to a pivotal role, with little redundancy, in maintaining root development and nutrient balance upon severe Zn deficiency.

## Conclusions

Altogether, this study showed that root growth inhibition upon Zn deficiency may be explained by a reduction of the final size of the mature cells, and a diminution of the number of dividing cells in the division zone of the RT, resulting from accelerated cell elongation and differentiation. We further showed that Zn allocation to the RT is prioritized upon Zn deficiency and identified ZIP12 as a key actor of the response to severe Zn deficiency. As such, this work provides a solid foundation for understanding the complex responses to Zn deficiency in the root tips of Arabidopsis.

## Acknowledgments

We thank Prof. Mark Aarts for the kind gift of pZIP4:GUS seeds, as well as A. Degueldre for technical support in ICP-OES analyses. We thank Prof. Stephan Clemens, Prof. Lieven de Veylder, Dr. Ana G.L. Assunção, and Dr. Grmay H. Lilay for helpful discussions. Funding was provided by the “Fonds de la Recherche Scientifique-FNRS” (CDR J.0009.17 to M.H.; PDR-T0120.18, PDR-T.0104.22 to M.H. and N.V.). The authors wish to thank the COST ACTION 19116 PLANTMETALS for efficient networking and discussion. No conflict of interest declared.

## Author contributions

MH and NV conceived and directed the research. NT, MH, NV and DPP designed the experiments. NT, DDP, PS, MS, MC, BB and SF performed all experiments. NT, DPP, NV and MH analysed the data. NT made the figures. NT, MH, MS and NV wrote the manuscript, with comments of all authors.

## Data availability

The RNA-Seq reads have been deposited in the National Center for Biotechnology Information (NCBI) Sequence Read Archive (SRA) Database with BioProject identification number (submission in progress). The other data that support the findings of this study are available from the corresponding authors upon reasonable request.

## Supporting Information

**Table S1.** Primers used for the mutant genotyping and RT-qPCR experiments.

**Table S2.** Expression of marker genes for the cell cycle, endocycle and DDR in the RNA-Seq analysis.

**Table S3.** Differentially expressed genes (DEGs) between remaining roots (RR) and root tip (RT) samples in control (1 µM Zn) and Zn deficiency (0 µM) conditions.

**Table S4.** Gene Ontology enrichment analysis amongst the differentially expressed genes (DEGs) less expressed in root tips (RT) than the remaining roots (RR) in control (1 µM Zn) and deficiency (0 µM) conditions.

**Table S5.** Gene Ontology enrichment analysis amongst the differentially expressed genes (DEG) more expressed in root tips (RT) than remaining roots (RR) in control (1 µM Zn) and deficiency (0 µM) conditions.

**Table S6.** Differentially expressed genes (DEGs) between control (1 µM Zn) and deficiency (0 µM) conditions in remaining roots (RR) and root tips (RT).

**Table S7**. Impact of Zn deficiency on developmental processes in root tips (RT) and remaining roots (RR).

**Supplementary Figure 1.** *zip12* mutant characterization.

**Supplementary Figure 2.** Comparison of cell elongation rate in the tip of the root in Arabidopsis plantlets grown control (1 µM Zn) or Zn deficiency (0 µM Zn) conditions.

**Supplementary Figure 3.** Cell cycle synchronization analysis.

**Supplementary Figure 4.** Illustration of methods.

**Supplementary Figure 5.** Impact of Zn deficiency on the expression of cell cycle phase, endocycle and DNA damage marker genes in Arabidopsis roots.

**Supplemental Figure S6.** Impact of Zn deficiency on root tip developmental and mitotic processes.

**Supplementary Figure 7.** Expression of *ZIP* and *NAS* genes in Arabidopsis roots at various Zn concentrations.

**Supplementary Figure 8.** Ionome profiling of the roots and shoots in Col-0 and *zip12-1* plantlets upon Zn deficiency.

**Methods S1.** This section presents detailed Materials and Methods.

