## Supporting Information for "Zinc deficiency induces spatially distinct responses in roots and impacts ZIP12-dependent zinc homeostasis in Arabidopsis"

The following Supporting Information is available for this article.

### Supporting tables

**Table S1.** Primers used for the mutant genotyping and RT-qPCR experiments.

**Table S2.** Expression of marker genes for the cell cycle, endocycle and DDR in the RNA-Seq analysis.

**Table S3.** Differentially expressed genes (DEGs) between remaining roots (RR) and root tip (RT) samples in control (1 µM Zn) and Zn deficiency (0 µM) conditions.

**Table S4.** Gene Ontology enrichment analysis amongst the differentially expressed genes (DEGs) less expressed in root tips (RT) than the remaining roots (RR) in control (1 µM Zn) and deficiency (0 µM) conditions.

**Table S5.** Gene Ontology enrichment analysis amongst the differentially expressed genes (DEG) more expressed in root tips (RT) than remaining roots (RR) in control (1 µM Zn) and deficiency (0 µM) conditions.

**Table S6.** Differentially expressed genes (DEGs) between control (1 µM Zn) and deficiency (0 µM) conditions in remaining roots (RR) and root tips (RT).

**Table S7**. Impact of Zn deficiency on developmental processes in root tips (RT) and remaining roots (RR).

### Supporting figures

**Supplementary Figure 1. *zip12* mutant characterization. a**. Position of the T-DNA insertion sites in the *zip12-1* and *zip12-2* mutants. **b-d**. Col-0, *zip12-1* and *zip12-2* plantlets were grown for two weeks on EDTA-washed agar plates in control (2 µM Zn) or Zn deficiency (0 µM Zn) conditions. **b**. *ZIP12* relative expression in Col-0, *zip12-1* and *zip12-2* plantlets. Values are means +/- SD from 3 technical replicates, relative to the mean expression into Col-0 roots in control condition (2µM Zn) . **c-d**. Primary root length of Col-0 and *zip12-2* (**c**) and *zip12-1* (**d**) in 2 µM Zn and 0 µM Zn. Different letters represent statistically different groups (ANOVA type II, with Tukey test correction, *p-value* < 0.05). Panel d is a fully independent experiment from, and confirms data presented in, **Fig. 7a**.

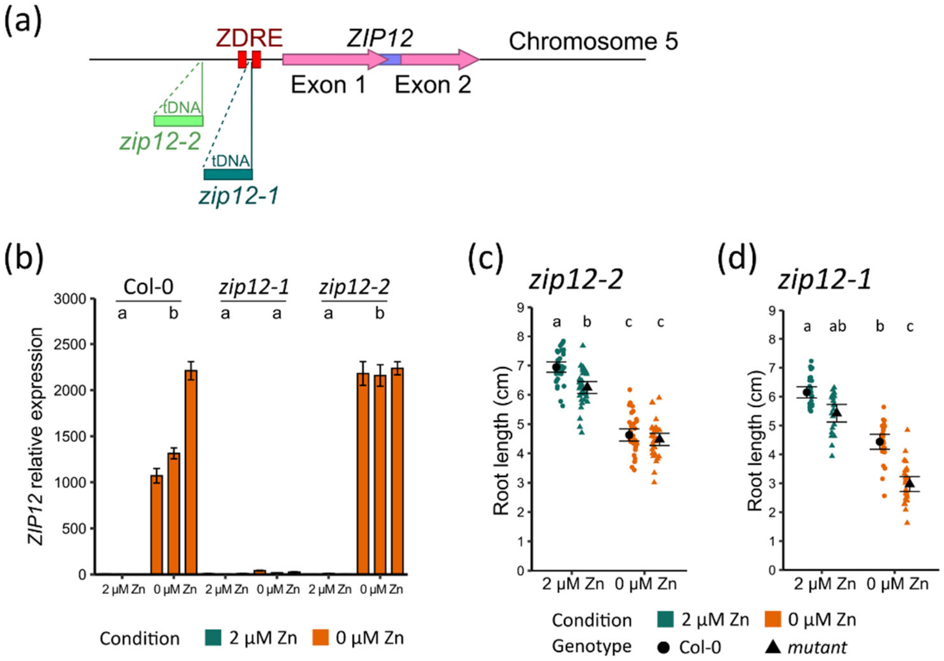

**Supplementary Figure 2**. **Comparison of cell elongation rate in the tip of the root in Arabidopsis plantlets grown control (1 µM Zn) or Zn deficiency (0 µM Zn) conditions.** Two linear regressions of the cortex cell length data from Fig. 2a were realized, (i) starting from the end of the RAM for 250 µm after the RAM (start of elongation zone) and (ii) from the beginning of the elongation zone to a distance of 1375 µm from the root tip (corresponding to the most distant data point for the 1 µM Zn condition). The regressions were realized in R with the linear model function lm(x) separately on the different zones and separately for the 1 µM Zn and 0 µM Zn data. The slopes of the regression lines provide an estimate of the cortex cell elongation rate in each zone.

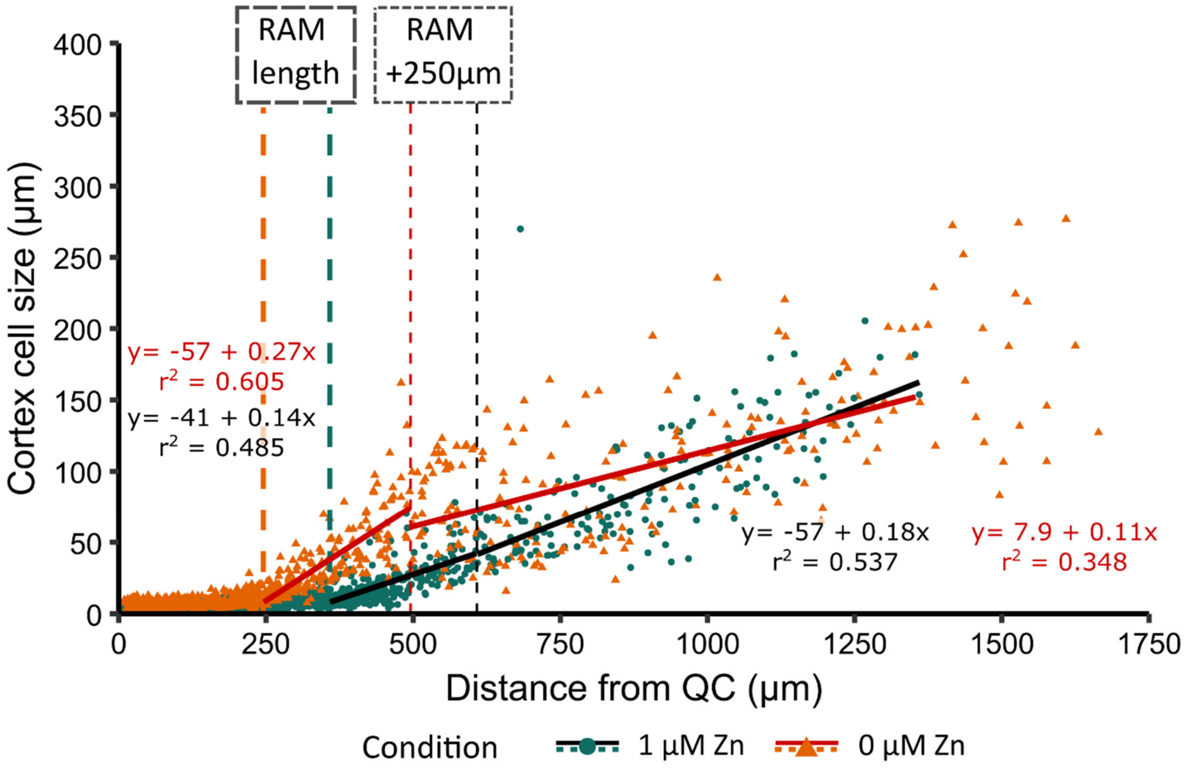

**Supplementary Figure 3. Cell cycle synchronization analysis.** Cell cycle was synchronized with 2 mM hydroxyurea in two-weeks old Arabidopsis plantlets grown in control conditions (1 µM Zn) or Zn deficiency (0 µM Zn). **a-b.** pCYCA3;1:CYCA3;1:GUS (**a**) and pCYCB1;2:CYCB1;2:GUS (**b**) plantlets were stained 0 h, 4 h, 8 h, 12 h, 16 h, 20 h, 22 h or 24 h, 26 h or 28 h after synchronization in control conditions (top) or Zn deficiency (bottom). **c**. GUS activity in pCYCB1;2:CYCB1;2:GUS was assessed by counting blue dots, corresponding to GUS-stained cells (Perilli & Sabatini, 2010; Thiébaut *et al.*, 2025), in the experiment represented in **b**. Stars correspond to significantly different values (ANOVA type I). The experiments were carried out on 5 plantlets per condition per time-point.

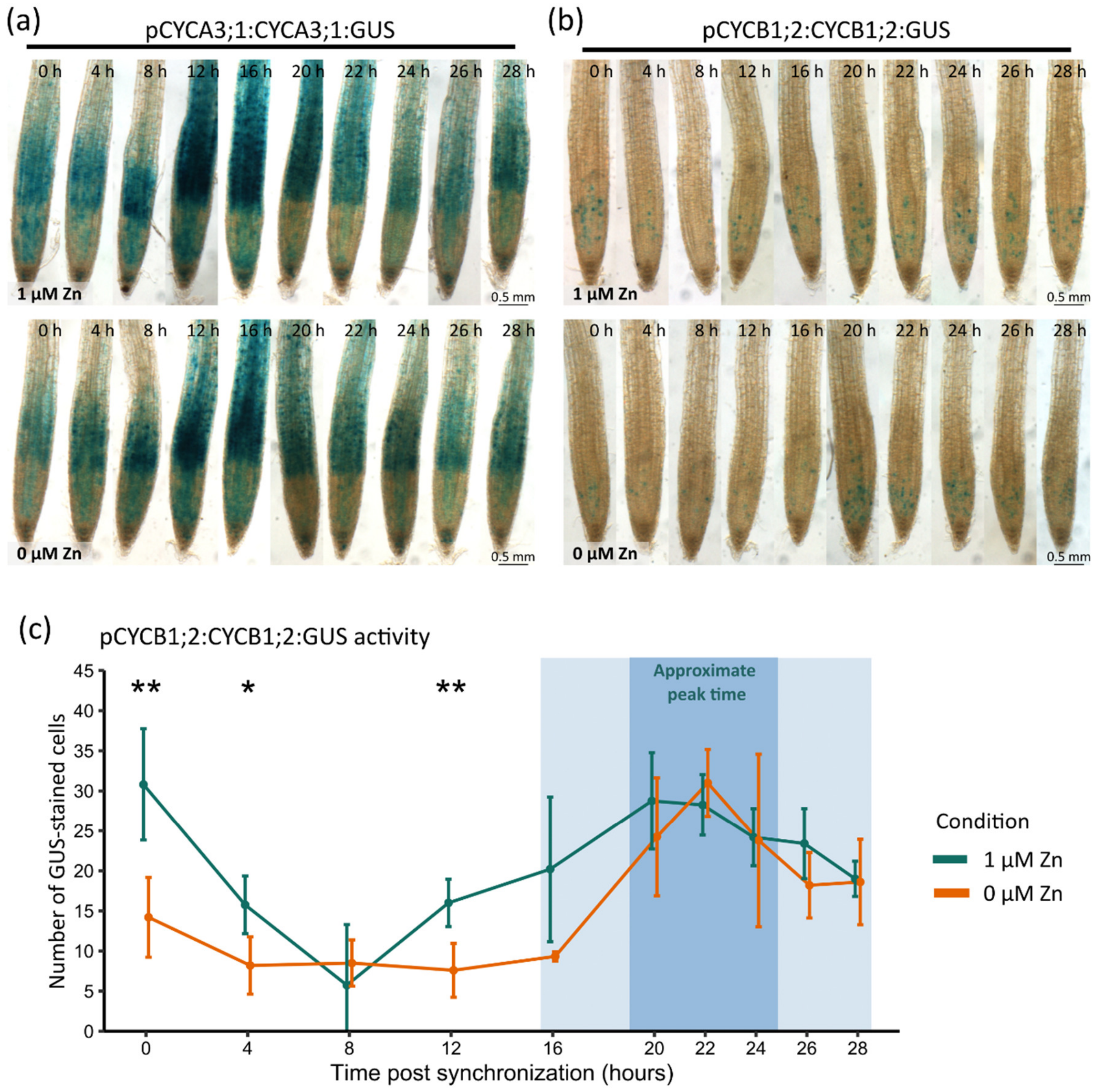

**Supplementary Figure 4. Illustration of methods. a**. Illustration of the measurement of cortex cell length and their distance from the quiescent centre. Propidium iodide-stained root apexes were mounted in water and imaged via confocal microscopy ($\lambda_{ex}$ = 543 nm and $\lambda_{em}$ within 600 and 730 nm). Scaled images were analysed using FIJI. Yellow lines represent the measurements of the cortex cell length and their distance from quiescent centre. **b-c**. Laser ablation ICP-MS: representative figure of dried samples and the corresponding potassium $^{39}K^{+}$ signal, used as a reference for confirming correct ablation, in root apical meristem (RAM) (**b**) and differentiated roots (**c**). In the $^{39}K^{+}$ signal panels, the green rectangles across the sections show how the data were collected for the graphs in **Fig. 4c-d,** as in (Thiébaut *et al.*, 2025).

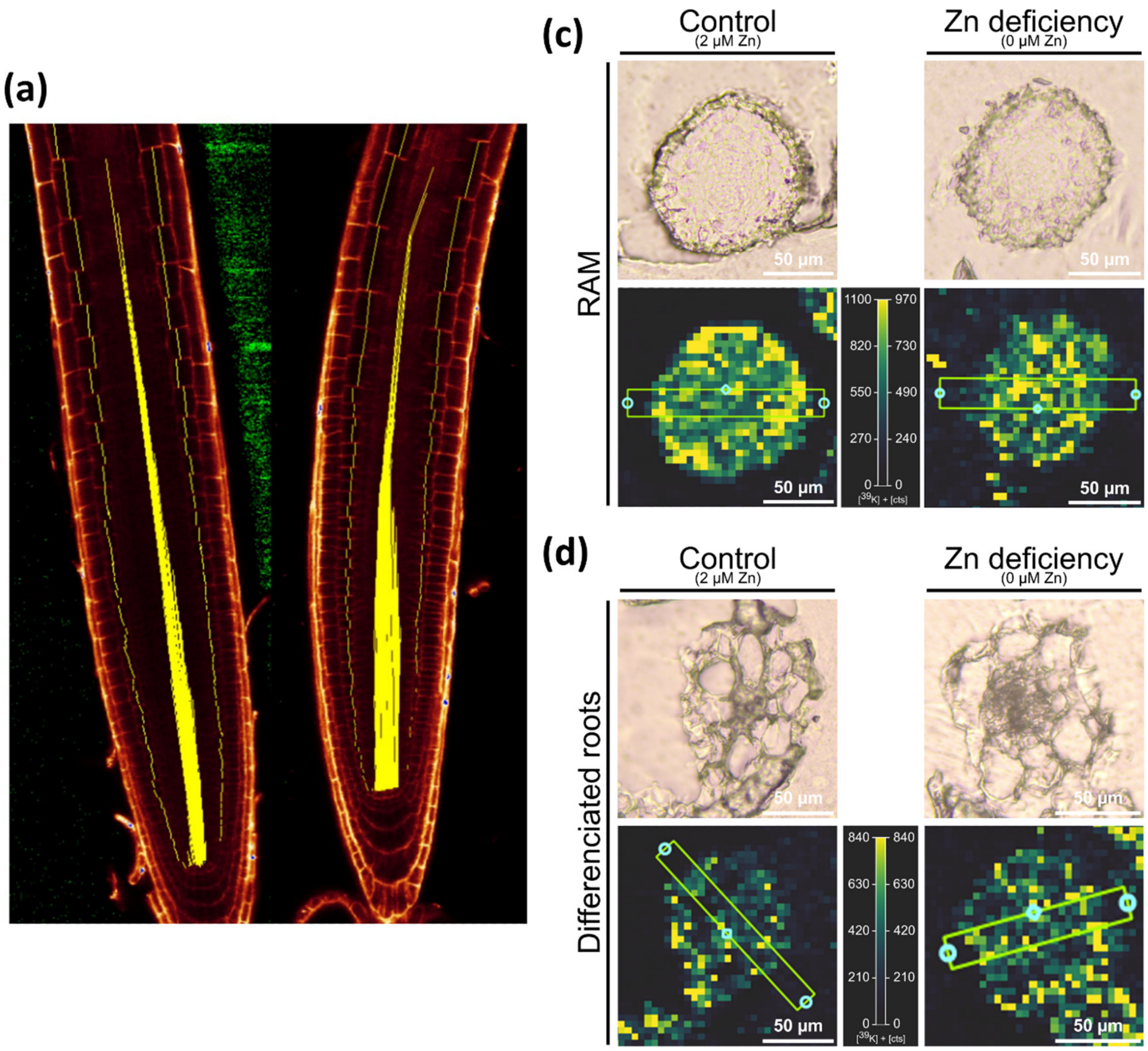

**Supplementary Figure 5. Impact of Zn deficiency on the expression of cell cycle phase, endocycle and DNA damage marker genes in Arabidopsis roots.** Arabidopsis plantlets were grown in three replicates for two weeks on EDTA-washed agar plates in control (1 µM Zn) or Zn deficiency (0 µM Zn) conditions. (a) Heatmap of $log_2$(fold change) of gene expression between Zn deficiency (0 µM Zn) and control conditions (1 µM Zn) in root tips (RT) and remaining roots (RR), based on RNA-Seq data. The expression of a manually curated list of 132 genes (**Table S2**) marking different phases of the cell cycle, endocycle and DNA Damage Response (DDR). (b) Genes differentially expressed (3/132) upon Zn deficiency in RR and RT. (c) Genes differentially expressed (9/132) in RT compared to RR in control or in Zn deficiency conditions. (b-c) * adjusted *p-value* < 0.05, n.s. not significant. upon Zn deficiency in RR and RT. d. Primary root growth inhibition of Col-0 and the *atm-1* mutant upon Zn deficiency. Coloured dots are 7 independent biological replicates, each dot representing the mean value from 18 plantlets. Black dots and whiskers represent mean values and standard deviations of all 7 experiments. The star shows statistically different groups (ANOVA type I, *p-value* < 0.05).

**(a)**

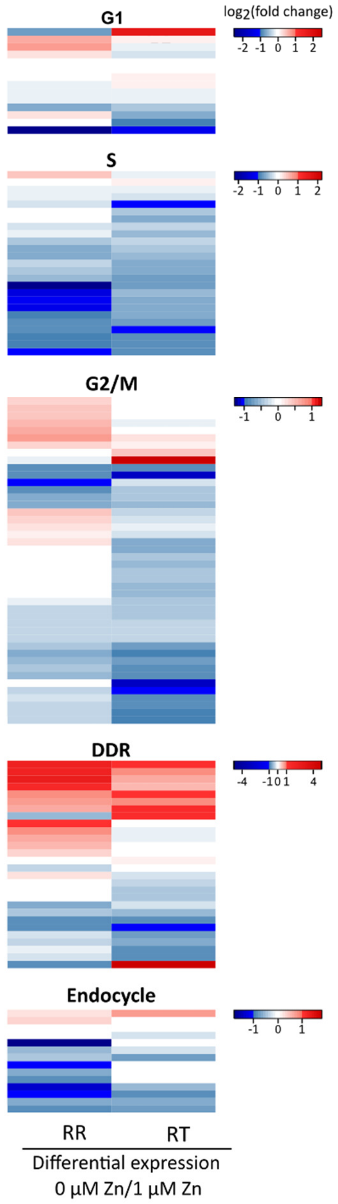


**(b)**

| Gene.ID | Gene | Phase | RR | RT |
|---|---|---|---|---|
| *AT1G19250* | *FMO1* | DDR | -0,84 $^{ns}$ | 4,93 * |
| *AT5G60250* | *AT5G60250* | DDR | 2,55 * | 0,68 $^{ns}$ |
| *AT5G10440* | *CYCD4;2* | G1 | -2,41 * | -1,24 $^{ns}$ |

**(c)**

| Gene.ID | Gene | Phase | 1 µM Zn | 0 µM Zn |
|---|---|---|---|---|
| *AT1G07500* | *SMR5* | DDR | -5,26 * | -4,50 * |
| *AT1G19250* | *FMO1* | DDR | -6,84 * | -1,04 $^{ns}$ |
| *AT2G18193* | *AT2G18193* | DDR | 2,56 * | 1,55 * |
| *AT5G60250* | *AT5G60250* | DDR | -0,60 $^{ns}$ | -2,43 * |
| *AT3G10525* | *SMR1* | Endocycle | 1,56 $^{ns}$ | 2,43 * |
| *AT4G22910* | *FZR2* | Endocycle | 1,32 $^{ns}$ | 1,99 * |
| *AT1G70210* | *CYCD1;1* | G1/S | -3,59 * | -1,07 $^{ns}$ |
| *AT3G50410* | *OBP1* | G1/S | -3,17 $^{ns}$ | -4,01 * |
| *AT4G34160* | *CYCD3;1* | G1/S | -1,02 $^{ns}$ | -1,97 * |

**(d)**

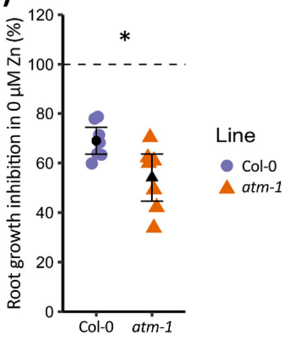

**Supplemental Figure S6. Impact of Zn deficiency on root tip developmental and mitotic processes.** Bulk RNA-Seq data from root tip (RT) and remaining root (RR) samples upon growth in control (1 µM Zn) or Zn deficiency (0 µM Zn) conditions were cross-referenced with the cell cycle phase-specific (**a**, Torii et al., 2020) or cell-type specific (**b**, Shahan et al., 2022) descriptions of gene expression in roots obtained by single-cell RNA-Seq. **a**. The figure represents the fraction of Zn deficiency DEG in either RR or RT (red and blue lines) matching to cell cycle scRNA-Seq clusters, as well as the size of the cell cycle clusters (grey bars) as defined by Torii *et al.* (2020). **b.** The figure represents the number of genes that were differentially expressed between RT and RR in control and Zn deficiency conditions, and that were among the 50 specific marker genes defined by Shahan *et al.* (2022) for each cell type and development stage.

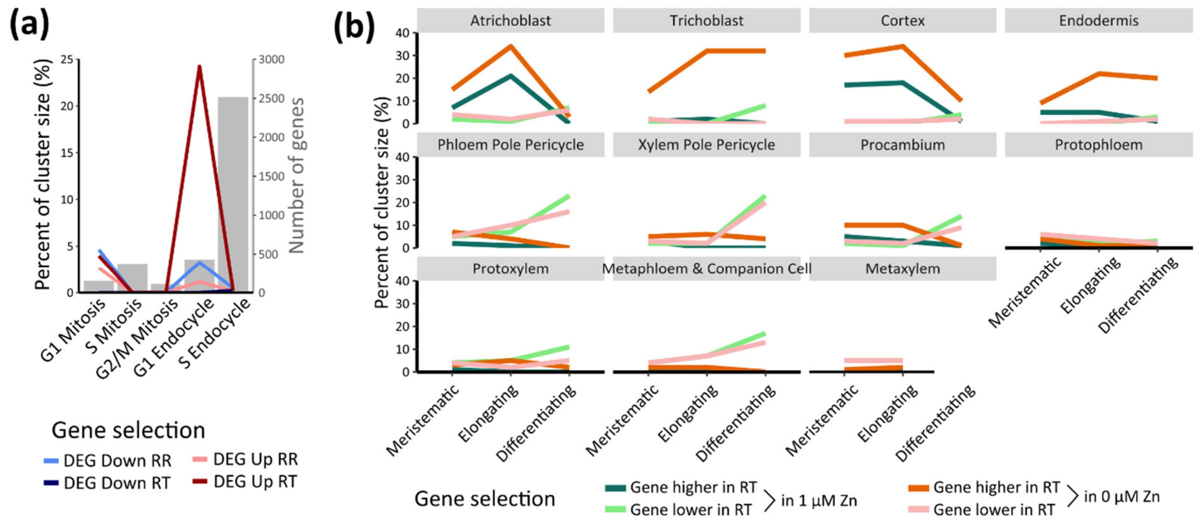

**Supplementary Figure 7. Expression of *ZIP* and *NAS* genes in Arabidopsis roots at various Zn concentrations.** Plantlets were grown for two weeks on EDTA-washed agar plates at various Zn concentrations (0-2 µM). **a-b.** Data from RNA-Seq analysis on plantlets grown in control (1 µM Zn) and Zn deficiency (0 µM Zn) conditions. **a.** Differential expression in $log_2$(fold change) upon Zn deficiency relative to 1 µM Zn in root tips (RT) and remaining roots (RR), adj. *p-val*: adjusted *p-value*, ns: non-significant. **b**. Distribution of gene expression [in $log_2$(counts)], for (i) all genes in RR samples (black line), (ii) the genes presented in panel (**a**) in the RR samples from plantlets grown in 1 µM Zn (green) or 0 µM Zn (orange). The vertical red line represents the 3$^{rd}$ quartile of the data composing the black line. **c.** *ZIP* gene expression in Col-0 plantlets assessed by RT-qPCR. Bar values and errors are means +/- SD from 3 technical replicates, relative to the mean expression in control condition (2 µM Zn). The RT-qPCR was realized in technical triplicates on three biological replicates (each including 180 plantlets).

**(a)**

| Gene.ID | Gene | $Log_2$(FC) $RR_{0\ \mu M\ vs\ 1\ \mu M}$ | $Log_2$(FC) $RT_{0\ \mu M\ vs\ 1\ \mu M}$ |
|---|---|---|---|
| *AT5G62160* | *ZIP12* | $7.3^{***}$ | $11.7^{***}$ |
| *AT1G56430* | *NAS4* | $1.2^{ns}$ | $2.2^{***}$ |
| *AT4G33020* | *ZIP9* | $1.1^{ns}$ | $3.3^{***}$ |
| *AT1G10970* | *ZIP4* | $0.85^{ns}$ | $1.9^{***}$ |
| *AT3G12750* | *ZIP1* | $0.85^{ns}$ | $2.2^{**}$ |
| *AT5G56080* | *NAS2* | $0.55^{ns}$ | $0.85^{ns}$ |
| *AT1G60960* | *IRT3* | $0.3^{ns}$ | $0.8^{ns}$ |
| *AT2G32270* | *ZIP3* | $0.1^{ns}$ | $1.6^{ns}$ |
| *AT1G31260* | *ZIP10* | $1.1^{ns}$ | $0^{ns}$ |

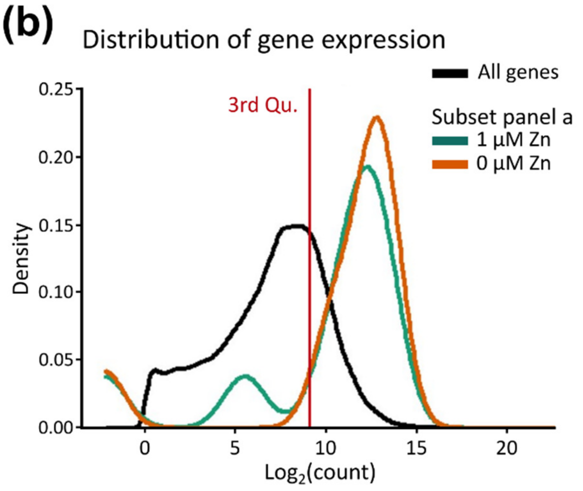


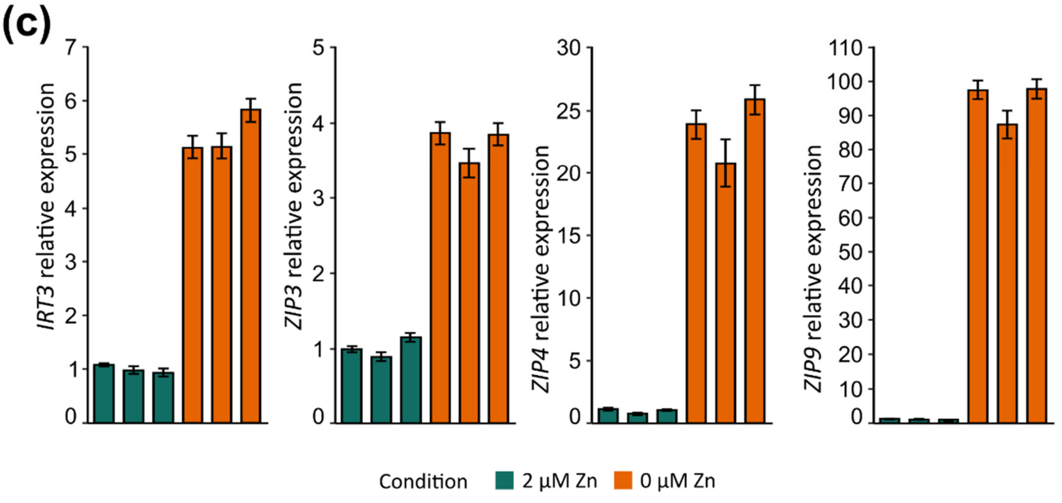

**Supplementary Figure 8. Ionome profiling of the roots and shoots in Col-0 and *zip12-1* plantlets upon Zn deficiency.** Col-0 and *zip12-1* plantlets were grown in control (2 µM Zn) or Zn deficiency (0 µM Zn) EDTA-washed agar plates. Zn, Cu and Mn concentrations are presented in **Fig. 7e**. Different letters represent statistically different groups (ANOVA type II, with Tukey test correction, *p-value* < 0.05).

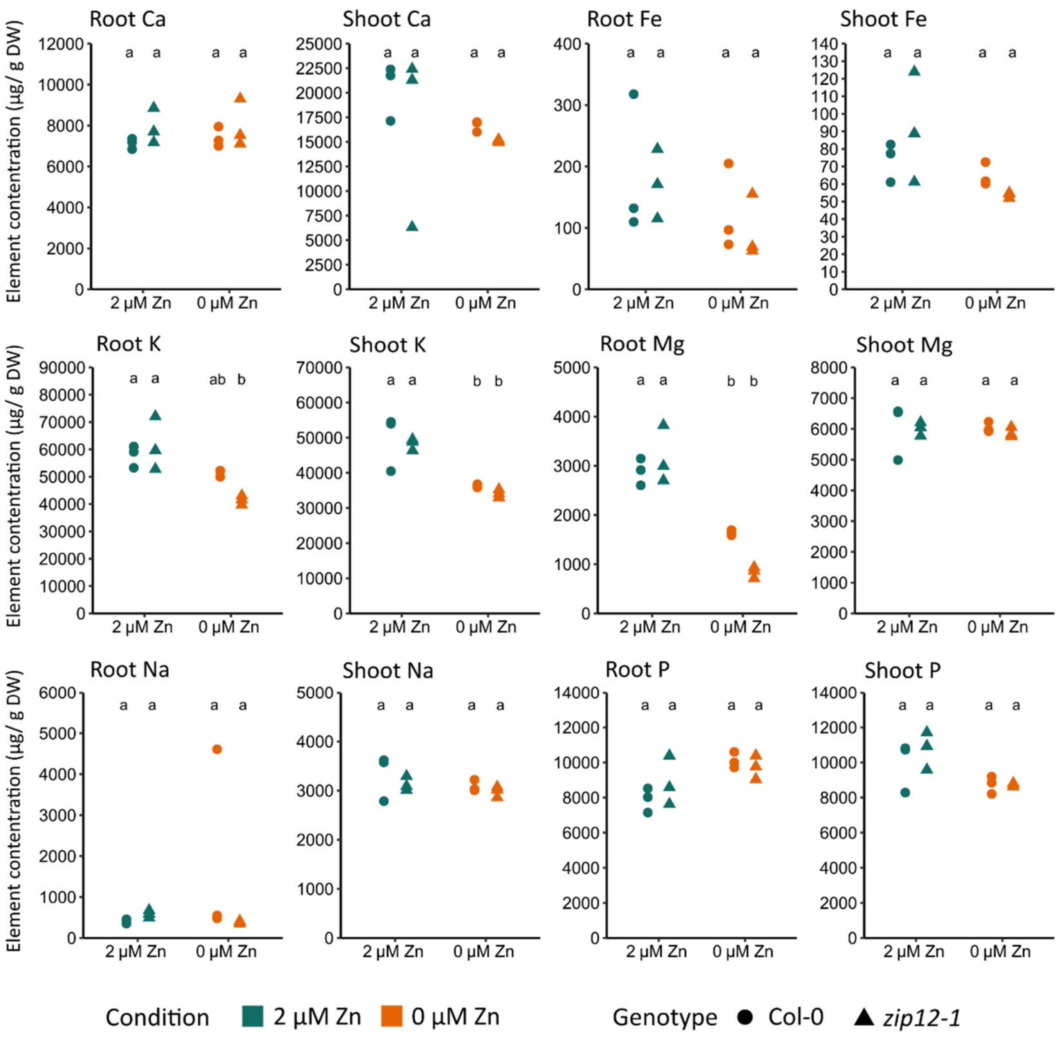

### Methods S1

This section presents detailed Materials and Methods.

#### Plant material, culture media and culture conditions

*Arabidopsis thaliana* (Col-0) was used in all experiments. The *ZIP12* (*AT5G62160*) mutants [*zip12-1* (SALK_137184), *zip12-2* (SALK_118705)] were obtained from NASC (Nottingham Arabidopsis Stock Centre, UK). These two mutants were genotyped using primers listed in **Table S1** to confirm T-DNA insertion sites (**Fig. S1**) and homozygous insertion lines were used in all experiments. The pWOX5:GUS, pZIP4:GUS (BG0011) and *atm-1* lines were previously described in Sarkar *et al.* (2007), Lin *et al.* 2016) and Garcia *et al.* (2003), respectively, and were kindly provided by Prof. L. De Veylder (Ghent University, Belgium) and Prof. M.G.M. Aarts (Wageningen, Netherlands). pCYCB1;2:CYCB1;2:GUS (N799897) and pCYCA3;1:CYCA3;1:GUS (N799893) lines ordered from NASC, have constructions encoding proteins that are subject to post-translational regulation through degradation (Kumpf *et al.*, 2014).

Seeds were surface-sterilized by NaClO/HCl gas exposure in a closed desiccator for 2h20 (Lindsey *et al.*, 2017). Seeds were then sown directly on 12x12 cm sterile petri plates with 50 mL of agar Hoagland medium (Charlier *et al.*, 2015) supplemented with 1 % sucrose (w/v) [1.5 mM $Ca(NO_3)_2$, 0.28 mM $KH_2PO_4$, 0.75 mM $MgSO_4$, 1.25 mM $KNO_3$, 0.5 µM $CuSO_4$, 1 µM $ZnSO_4$, 5 µM $MnSO_4$, 25 µM $H_3BO_3$, 0.1 µM $Na_2MoO_4$, 50 µM KCl, 10 µM Fe-HBED, 3 mM MES-KOH pH 5.7] and kept in the dark for 2 days at 4°C. Plates were incubated vertically for 2 weeks at 20°C in a short-day cycle (8h light, 100 $\mu mol.m^{-2}.s^{-1}$ / 16h dark) in Grobank climate-controlled growth chamber (CLF Climatics, Germany). For laser-ablation ICP-MS, culture conditions were slightly different with cycles of 8 h light (21°C), 150 $\mu mol.m^{-2}.s^{-1}$ / 16h dark (19°C).

To achieve Zn deficiency, Zn trace-contaminations were limited as much as possible as follows. Each glass vial containing agar, media and stock solutions was rinsed once with HCl 6 M, then 7 times with $ddH_2O$ prior use in order to remove the micronutrient contaminations from glass. The agar (Type M, Sigma-Aldrich, USA) was EDTA-washed as follows: 12 g/L agar were weighted, transferred in a 2L acid-washed Erlen-Meyer together with 250 mL of EDTA 5 mM (pH 8, with KOH) and agitated on an orbital shaker [70 rotations per minute (RPM)] for 5h. Then, 1h rest was observed for agar sedimentation, 35 mL of supernatant were discarded and $ddH_2O$ was added up to 800 mL. The rinsing procedure was repeated 7 times, twice a day. Hoagland medium supplemented with 1 % sucrose (w/v) was finally prepared in the washed agar-containing Erlen-Meyer. $ZnSO_4$ was added in various concentrations (0 µM, 1 µM, 2 µM or 5 µM) as indicated. Unless otherwise stated, all experiments were conducted in three independent replicates. The replication level of each experiment is detailed in Figure legends.

#### Root growth and RAM phenotyping

Plates containing the two week-old plantlets were scanned on a Epson perfection scanner, and primary roots were measured using FIJI (https://imagej.net/Fiji). To measure the

Root Apical Meristem (RAM) size, two weeks-old plantlets were rinsed 30 seconds in distilled water, then stained for 2 minutes in Propidium iodide (33.4 mg/L) and rinsed again in distilled water. Roots were mounted on glass slides and visualized with a SP2 TCS AOBS confocal laser-scanning inverted microscope (Leica, Mannheim, Germany) with excitation at 543 nm and emission taken between 600 and 730 nm. The pictures of the propidium iodide-stained roots were then used to quantify several morphological features using FIJI, as follows. The length of the meristematic zone was assessed as the distance between the first elongated cortex cell and the quiescent centre (QC) [(Perilli & Sabatini, 2010), **Fig. S4**]. The number of cortex cells in this zone was simultaneously assessed and a mean cortex cell size in the RAM was calculated by dividing RAM length (in µm) by the number of cortex cells for each root. Additionally, individual cortex cell size and their specific distance from the QC were measured to represent the elongation of the cells within the first mm of the root tips (**Fig. S4**). To measure the size of the elongation zone, the distance between the first elongated cortex cell and the first root hair was measured for each root. Finally, a picture at 5-6 mm from the root tip, in an area where root hairs are present, was used to measure the length of 5 cortex cells per root. Cell death, visible with propidium iodide staining inside the cell, was also noted if observed.

**GUS staining and MUG assay**

Two-week-old plantlets were harvested in cold acetone (90 % v/v) and then incubated at room temperature for 20 min. Plantlets were rinsed in staining buffer [0.2 % Triton X-100, 2 mM $Fe(CN)_6^{3-}$ (Ferrocyanide), 2 mM $Fe(CN)_6^{4-}$ (Ferricyanide), 7.7 mM $NaH_2PO_4$ and 17.1 mM $Na_2HPO_4$] and infiltrated with staining buffer in the presence of 5-bromo-4-chloro-3-indolyl-β-D-glucuronide (104.4 µg/mL, X-Gluc) by applying vacuum until sinking of the plantlets. The staining reaction was then conducted at 37°C (25 min for pZIP4:GUS, 15 min for pWOX5:GUS, 1h15 for pCYCA3;1CYCA3;1:GUS and 1h45 for pCYCB1;2:CYCB1;2:GUS). The staining reaction was stopped by a 30 min incubation in EtOH 30 % at room temperature, followed by 30 min incubation in FAA [Formaldehyde 5 % (v/v), Acetic acid 10 % (v/v), EtOH 50 % (v/v)] and finally overnight at 4°C in EtOH 70 % before imaging with a Nikon SMZ1500 stereomicroscope equipped with a Nikon Digital Sight DS-5M camera.

To perform cell cycle synchronization, two week-old plantlets were transferred onto fresh solid 1 µM or 0 µM Zn Hoagland media supplemented with 2 mM hydroxyurea, and quickly placed back in controlled culture conditions (Cools *et al.*, 2010). At each time point, 5 plantlets of pCYCA3;1CYCA3;1:GUS and pCYCB1;2:CYCB1;2:GUS were harvested in cold acetone and GUS staining was performed as described above.

Quantification of GUS activity was conducted differently for the different reporter lines. Being very weak, it was quantified by counting the blue spots (Perilli & Sabatini, 2010) in GUS-stained pCYCB1;2:DB-CYCB1;2:GUS lines. To do so, microscope pictures were acquired serially across the root depth to account for all RAM cell layers, and cells were counted with FIJI cell counter Plugin (**Fig. S4**). Being much stronger, GUS activity was measured via MUG (4-Methylumbelliferyl-Glucuronide) assay in pCYCA3;1:CYCA3;1:GUS lines as follows. Approximately 120 root tips per sample (~3 mm length, **Fig. S3**) were harvested in liquid nitrogen and grinded twice 1 min with a mixer mill (MM200, Retsch, Germany) at 25 Hz in the presence of steel beads. It was then re-suspended in protein extraction buffer [50 mM Na-phosphate buffer (pH 7.4), 10 mM EDTA, 0.1 % (w/v) SDS, 0.1 % (v/v) Triton X-100 and 10 mM β-mercaptoethanol] and grinded once more 1 min at 25 Hz. Cell debris were separated by centrifugation (1 min, 15 000 *g*, 4°C) to obtain a protein extract in the supernatant. Protein quantity in the extract was determined with a Bradford assay (Bio-Rad, USA), against a BSA standard curve. For activity measurement, 20 µL of protein extract was incubated in 150 µL of extraction buffer in the presence of 2 mM MUG. The reaction was stopped after 3 h with 150 µL $Na_2CO_3$ (200 mM) and the fluorescence emitted by the released methylumbelliferone (MU) was measured with a VICTOR Nivo (PerkinElmer, USA) plate reader, with an excitation at 355 nm and absorption measured with a 460/30 nm filter. MU quantity was assessed by comparison to standard curve and activity was calculated in pmol MU $min^{-1}$ mg $protein^{-1}$.

**RNA preparation**

For RNA-Seq analysis, each RT sample (~2 mm length) was collected and pooled from 36 square petri dishes each containing 18 Col-0 plantlets (~650 plantlets) per condition: the RTs were directly cut on the agar using clean microdissection scissors (Stainless model SI-50-4500, Schreiber, Germany) (Thiébaut *et al.*, 2025), quickly pooled in a RNAse free microcentrifuge tube, and kept frozen in liquid nitrogen. Remaining roots (RR) was pooled in a second associated sample. The experiment was conducted in biological triplicates, generating a total of 12 samples [RT and RR in 2 conditions (0 µM Zn and 1 µM Zn)]. Total RNAs were extracted with the Maxwell® RSC Plant RNA Kit (including a DNAse step, Promega, USA) using a Maxwell® RSC robot (Promega, USA). Total RNA quality was controlled on a Bioanalyzer using a QC RNA kit (Agilent, USA) before library preparation.

For reverse transcription quantitative polymerase chain reaction (RT-qPCR), samples were collected in biological triplicates. Each of the triplicate contained the material from several petri dishes: 9 square petri dishes each containing 18 plantlets per sample, in two conditions (0 µM Zn and 2 µM Zn) and grinded with tungsten beads 2x45 seconds using a mixer mill at 25 Hz. Total RNAs were extracted using the RNAeasy Microkit (including a DNAse step, Qiagen, Germany).

**RT-qPCR**

cDNAs were prepared from 300 ng total RNAs using Oligo(dT) with the RevertAid H Minus First Strand cDNA Synthesis Kit (Fisher Scientific, USA). RT-qPCR was conducted in technical triplicates for each 15x-diluted cDNA sample/primer combinations with the Takyon Low Rox SYBR MasterMix dTTP Blue (Eurogentec, Belgium) kit according to the manufacturer's protocol, on a QuantStudio5 thermocycler (Applied Biosystems, Thermofischer, USA) with the primer listed in **Table S1**. Primer pair efficiency was calculated with LinReg PCR software (Ramakers *et al.*, 2003) and expression normalized with the expression of three housekeeping genes [*At5g60390-EF1alpha*, *At4g05320-UBQ10*, and *At1g58050,* (Nouet *et al.*, 2015)] and relative to the control (2 µM Zn) in the qBase+ software (Biogazelle, Belgium).

### RNA-Seq, library preparation, sequencing, data analysis and representation

Library preparation was performed from 500 ng total RNAs using the TruSeq Stranded mRNA Sample Preparation Kit (Illumina, USA). Library quality was assessed with a QIAxcel screening kit (Qiagen, Germany). Quantification, prior dilution and pooling of the libraries was done using the KAPA SYBR® FAST Universal qPCR Kit and an Applied Biosystems 7900HT Real-Time PCR system. RNA-Sequencing was performed on a NovaSeq6000 at the GIGA Genomics platform of ULiège (flowcell S4 V1.5, with 2x paired-end reads of 150 nucleotides) to obtain an average of 21.3 million reads per sample. Upon quality control using FastQC (v0.10.1, http://www.bioinformatics.babraham.ac. Uk/projects/fastqc/), quality filtering was conducted using Trimmomatic (v0.32, Bolger *et al.*, 2014) with the following parameters (leading = 30, trailing =30, slidingwindow = 10:30, crop = 98, minlen = 98) and Prinseq (dust = 90, https://prinseq.sourceforge.net/). After quality filtering, an average of 17.4 million reads/sample were conserved. HiSAT2 (Kim *et al.*, 2019) was used to map reads on the TAIR10 Arabidopsis genome assembly version with Araport11 annotation (Cheng *et al.*, 2017). An average of ~14.4 million reads were aligned once per sample. Raw read counts per gene per sample were then obtained using htseq-count (v0.6.1p1,Anders *et al.*, 2015).

Read counts and sample description datasets were analysed with DESeq2 package in R. Differential analysis was conducted with the function DESeq2::results() with the parameters lfcThreshold = 1, alpha = 0.05, followed by a selection of padj < 0.05 and either log2FoldChange > 1 or log2FoldChange < -1 for up-regulated and down-regulated genes, respectively. Read counts were normalized with the function DESeq2::rlog(). Representations were done using the following functions: DESeq2::plotPCA(), DESeq2::plotcount() and gplots::heatmap.2() on as.matrix(dist(t(assay(rld)))).

A manually assembled list of cell cycle marker genes used for analysis is presented in **Tables S4, S5**. GO enrichment analysis for biological processes was conducted using the Panther Database (https://pantherdb.org) in September 2022, against *A. thaliana* annotated genes, with Bonferroni-correction with *p-value* < 0.05.

Identification of the transcription factors was done by comparing DEG lists to the *A. thaliana* Transcription Factor DataBase (AtTFDB) from the Arabidopsis Gene Regulatory Information Server (AGRIS, https://agris-knowledgebase.org/AtTFDB/).

### Mineral analysis

Upon growth in control or Zn deficiency conditions in Hoagland agar plates, shoots and roots of 160-180 two-week-old plantlets per sample were separated. Pooled shoots were rinsed in 4°C ultrapure Milli-Q water (Millipore) and dried at 55°C. Pooled roots were desorbed twice 10 minutes in $CaCl_2$ 5 mM, MES 1 mM (pH 5.7) at 4°C and rinsed twice 5 min in ultrapure Milli-Q water (Millipore, USA) and dried at 55°C. Dried root and shoot samples were weighted and digested in 3 mL $HNO_3$ (65 %) by gradual heating (15 min at 45°C, 15 min at 65°C) until 95°C for a 60 min incubation using a DigiPrep Graphite Block Digestion System (SCP Science, AnalytiChem, Canada). Digested samples were diluted to a final volume of 10 mL before ICP-

OES (Inductively Coupled Plasma Optical Emission Spectroscopy) using a 5110 SVDV ICP-OES device (Agilent, USA).

**Zinc imaging by Laser Ablation ICP-MS**

Upon growth for two weeks on EDTA-washed agar Hoagland plates in control (2 µM Zn) and Zn deficiency (0 µM Zn) conditions, roots were sampled as 12 µm-thick sections, from two regions of interest (ROI): the RAM (at ~200 µm from the tip of the columella) and differentiated roots (at a distance from the root apex corresponding to 50-60 % of the total root length).

At harvest time, ROI were aligned in a "root bouquet" structure, prepared and sectioned as described (Thiébaut *et al.*, 2025). Fourteen-µm thick sections were dried and photographed and selected for Laser Ablation-ICP-MS.

The Laser Ablation-ICP-MS system was composed of an Iridia laser ablation system (Teledyne inc., UK), coupled to a triple quadrupole ICP-MS [ICP-QQQ-MS 8900 (Agilent, UK)]. Samples were ablated at a spot size of 5 µm diameter, at 200 Hz, with a fluence of 0.5 $J.cm^{-2}$. For the detection, two elements were analysed: $^{39}K$ and $^{66}Zn$ with integration times of 0.001 and 0.07 s, respectively. The HDIP software was used for data analysis (Teledyne CETAC Technologies, USA).
